# NADPH composite index analysis quantifies the relationship between compartmentalized NADPH dynamics and growth rates in cancer cells

**DOI:** 10.1101/2024.04.27.591477

**Authors:** Sun Jin Moon, Anush Chiappino Pepe, Wentao Dong, Joanne K. Kelleher, Matthew G. Vander Heiden, Gregory N. Stephanopoulos, Hadley D. Sikes

## Abstract

NADPH, a highly compartmentalized electron donor in mammalian cells, plays essential roles in cell metabolism. However, little is known about how cytosolic and mitochondrial NADPH dynamics relate to cancer cell growth rates in response to varying nutrient conditions. To address this issue, we present NADPH composite index analysis, which quantifies the relationship between compartmentalized NADPH dynamics and growth rates using genetically encoded NADPH sensors, automated image analysis pipeline, and correlation analysis. Through this analysis, we demonstrated that compartmentalized NADPH dynamics patterns were cancer cell-type dependent. Specifically, cytosolic and mitochondrial NADPH dynamics of MDA-MB-231 decreased in response to serine deprivation, while those of HCT-116 increased in response to serine or glutamine deprivation. Furthermore, by introducing a fractional contribution parameter, we correlated cytosolic and mitochondrial NADPH dynamics to growth rates. Using this parameter, we identified cancer cell lines whose growth rates were selectively inhibited by targeting cytosolic or mitochondrial NADPH metabolism. Mechanistically, we identified citrate transporter as a key mitochondrial transporter that maintains compartmentalized NADPH dynamics and growth rates. Altogether, our results present a significant advance in quantifying the relationship between compartmentalized NADPH dynamics and cancer cell growth rates, highlighting a potential of targeting compartmentalized NADPH metabolism for selective cancer cell growth inhibitions.

## Introduction

Reduced nicotinamide adenine dinucleotide phosphate (NADPH) is an essential molecule in living organisms by virtue of its function as an electron donor that supports reductive biosynthesis and protects cells against oxidative stress^1,2^. In eukaryotic cells, the inner mitochondrial membranes is considered to be impermeable to NADPH, leading to separate cytosolic and mitochondrial pools of this cofactor^3^. Widely used methods exist that are both robust and sensitive to assess NADPH levels in cells, but measuring subcellular concentration of NADPH is challenging because it can be become oxidized during the cell lysis or fractionation steps that are required^4^. Other methods such as deuterium-labeled isotope tracing approaches, fluorescent protein biosensors, or genome-scale metabolic network analysis have been used to reveal compartmentalized NADPH metabolism, but an ability to assess levels and dynamics across multiple compartments is lacking^5–10^. Recently, a genetically encoded sensor for NADPH, called iNap, was developed to directly measure compartmentalized NADPH levels and its dynamics in live, single cells^11^. The sensor was used to reveal the response of cytosolic NADPH dynamics to oxidative stress in glucose-fed or deprived conditions^11^, and the response of cytosolic and mitochondrial NADPH dynamics to mitochondrial oxidative stress generated by D-amino acid oxidase system^12^. Yet, iNap sensors have not been applied to evaluate compartmentalized NADPH dynamics in diverse nutrient microenvironment.

Nutrient availability in tumor microenvironment plays an important role in NADPH metabolism as it can impact cytosolic and mitochondrial NADPH pools^13^. For instance, glucose, glutamine, and serine have been considered as key nutrients whose metabolism supports regeneration of NADPH in the cytosol and, or mitochondria^14,15^. Glucose metabolism can produce NADPH through the pentose phosphate (PP) pathway^16^, and cancer cells have been reported to upregulate the PP pathway to supply NADPH for fatty acid synthesis and to combat reactive oxygen species stress^17^. Withdrawing glucose diminishes the cytosolic NADPH pool, making cells vulnerable to oxidative stress^11^. The contributions of serine and glutamine metabolism to compartmentalized NADPH levels depends on the location of specific isoforms of NADPH generating enzymes such as malic enzyme (ME), isocitrate dehydrogenase (IDH), or methylenetetrahydrofolate dehydrogenase (MTHFD)^5,14,15^. Knockout of cytosolic or mitochondrial isoforms of IDH, ME, or MTHFD can all decrease cellular NADPH/NADP^+14,15,18,19^, but how this affected compartmentalized cofactor levels is not known. Genome-scale metabolic models also predicted knockout of IDH or ME reduced the net generation cellular NADPH in head and neck squamous cell carcinoma^8^; however, testing these models as well as how changes in extracellular nutrient substrates impact compartmentalized NADPH levels is less clear.

Moreover, while cytosolic and mitochondrial NADPH pools are separately maintained by organelle specific NADPH producing enzymes, NADPH reducing equivalent can be transferred between compartments via indirect metabolite shuttle systems^20,21^. Several metabolite shuttle systems involve metabolites pairs such as malate-pyruvate, (iso)citrate-αKG, citrate-malate-pyruvate, and glutamate-malate, that enable transfer of NADPH between the cytosol and mitochondria^20^. Isotope tracing and metabolomics revealed that citrate transporter, encoded by *SLC25A1*, mediates transfer of NADPH from cytosol to mitochondria in order to suppress oxidative stress during anchorage-independent growth^22^, while the dicarboxylate carrier, encoded by *SLC25A10*, contributes to the indirect transfer of NADPH between mitochondria and cytosol^23^. Recently, *SLC25A51* was shown to involve in transferring NAD^+^ to mitochondria^24^, raising the possibility that direct NADP(H) transporters may exist. Currently, most efforts to understand compartmentalized NADP(H) transport have focused on individual transporters, lacking a systematic evaluation of the mitochondrial transporters that play significant role in maintenance of compartmentalized NADPH redox states.

Here, we report NDAPH composite index analysis, which combines iNap sensors, a high-throughput image analysis pipeline, and a correlation analysis, to quantify cytosolic and mitochondrial NADPH dynamics in response to various nutrient conditions and establish their relationship with growth rates. We not only find the distinct NADPH dynamics across the cancer cell lines, but also determine cell-line specific fractional contribution parameter, which estimates a relative contribution of cytosolic and mitochondrial NADPH to growth rates. Using this parameter as a guidance, we demonstrate selective growth inhibition in cancer cell lines, such as MDA-MB-231 and HCT-116, by perturbing cytosolic or mitochondrial NADPH metabolism, respectively. Moreover, using a reconstructed genome-scale metabolic model (GEM) for gene-knockout simulation and flux analysis^25,26^, we systematically evaluate the *SLC25* mitochondrial transporter gene family, identify and experimentally validate that *SLC25A1* plays a key role in maintaining both cytosolic and mitochondrial NADPH pools in shuttling NADPH. Given the essential role of compartmentalized NADPH metabolism in cell growth and survival, targeting NADPH metabolism has been proposed as a potential anti-cancer therapy^27,28^. To this end, our NADPH composite index analysis quantifies the relationship between cell-type dependent compartmentalized NADPH dynamics and growth rates, demonstrating its potential to serve as a guide to target compartmentalized NADPH metabolism for selective cancer cell growth inhibitions.

## Results

### NADPH composite index analysis quantifies cytosolic and mitochondrial NADPH dynamics

Here, we present NADPH composite index analysis, which quantifies the relationship between compartmentalized NADPH dynamics and growth rates in response to varying nutrient conditions using a fractional contribution parameter (F). The optimal F values serves as a guide to target cytosolic or mitochondrial NADPH metabolism for selective growth inhibitions of cancer cells **(Fig. 1)**. The analysis mainly consists of three steps: 1) preparation of NADPH sensors and an automated image analysis pipeline, 2) assessment of compartmentalized NADPH dynamics in response to varying nutrients conditions such as glucose, glutamine, and serine deprivation, and 3) determination of the optimal F value to selectively inhibit cancer cell growths by targeting either cytosolic or mitochondrial NADPH metabolism (***SI Appendix*, Fig. 1*A*).** First, as a tool to measure cytosolic and mitochondrial NADPH, we used iNap sensor, a fluorescent protein-based biosensor whose fluorescence spectrum changes upon variation of NADPH levels and can be expressed in mitochondria using a mitochondrial localization tag (***SI Appendix*, Fig. 1*B***)^11^. Since the circularly permuted yellow fluorescent protein used in iNap sensor is sensitive to the change of pH upon excitation at 488 nm, we created control cell lines that stably expressed iNap-ctr sensors, which lack NADPH binding affinity and respond only to pH changes. Based on these measurements, we defined NADPH index, which is a pH-corrected sensor output by normalizing iNap fluorescence ratio to the iNap-ctr fluorescence ratio (***SI Appendix*, Fig. 1*C*)**.

**Figure 1.**
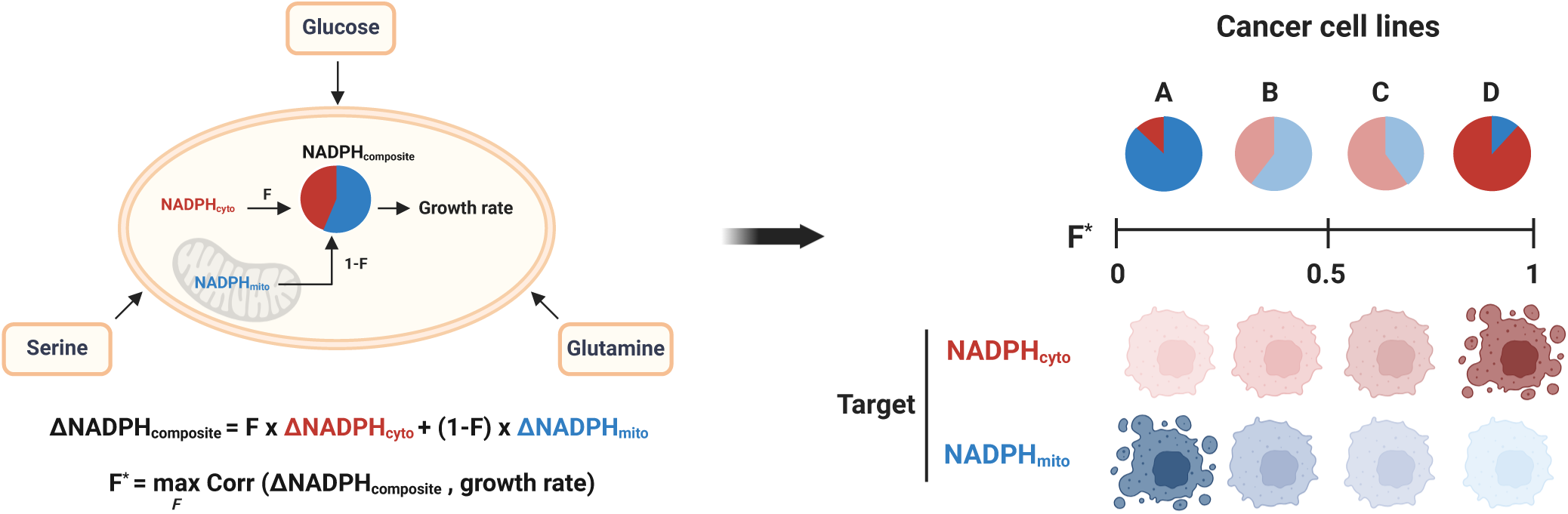
Overview of NADPH composite index analysis. NADPH composite index analysis quantifies the relationship between cytosolic and mitochondrial NADPH dynamics with cancer cell growth rates in response to varying nutrients conditions, using a a fractional contribution parameter (F). The optimal F value (F*) serves as a guide to target cytosolic or mitochondrial NADPH metabolism for selective growth inhibitions of cancer cells. Growth rates of cancer cell lines with F* closer to 1 are sensitive to the perturbation of cytosolic NADPH metabolism, whereas those with F* closer to 0 are sensitive to the perturbation of mitochondrial NADPH metabolism.

Next, as manually measuring the compartmentalized NADPH indices from fluorescence images is low-throughput, we designed a pipeline that used CellProfiler^29^ with optimally determined parameter settings to process thousands of images in a high-throughput manner. To test this pipeline, we generated Hela cell lines that stably expressed cytosolic or mitochondrial iNap sensors and assessed compartmentalized NADPH dynamics under glucose deprived conditions (**Fig. 2*A***). Our pipeline with optimized parameters identified individual cells that expressed cyto- or mito-iNap sensors as individual objects in each condition (**Fig. 2*B***). In the absence of glucose, we observed that both iNap (red) and mito-iNap (blue) fluorescence intensity at 488 nm increased but the intensity at 415 nm decreased compared to the control, a condition in which the cells were cultured in presence of glucose (black) **(Fig. 2*C*)**. The corresponding fluorescence ratio (F_415nm_/F_488nm_) of iNap and mito-iNap decreased compared to the control, suggesting a decrease of cytosolic and mitochondrial NADPH level **(Fig. 2*D*)**.

**Figure 2.**
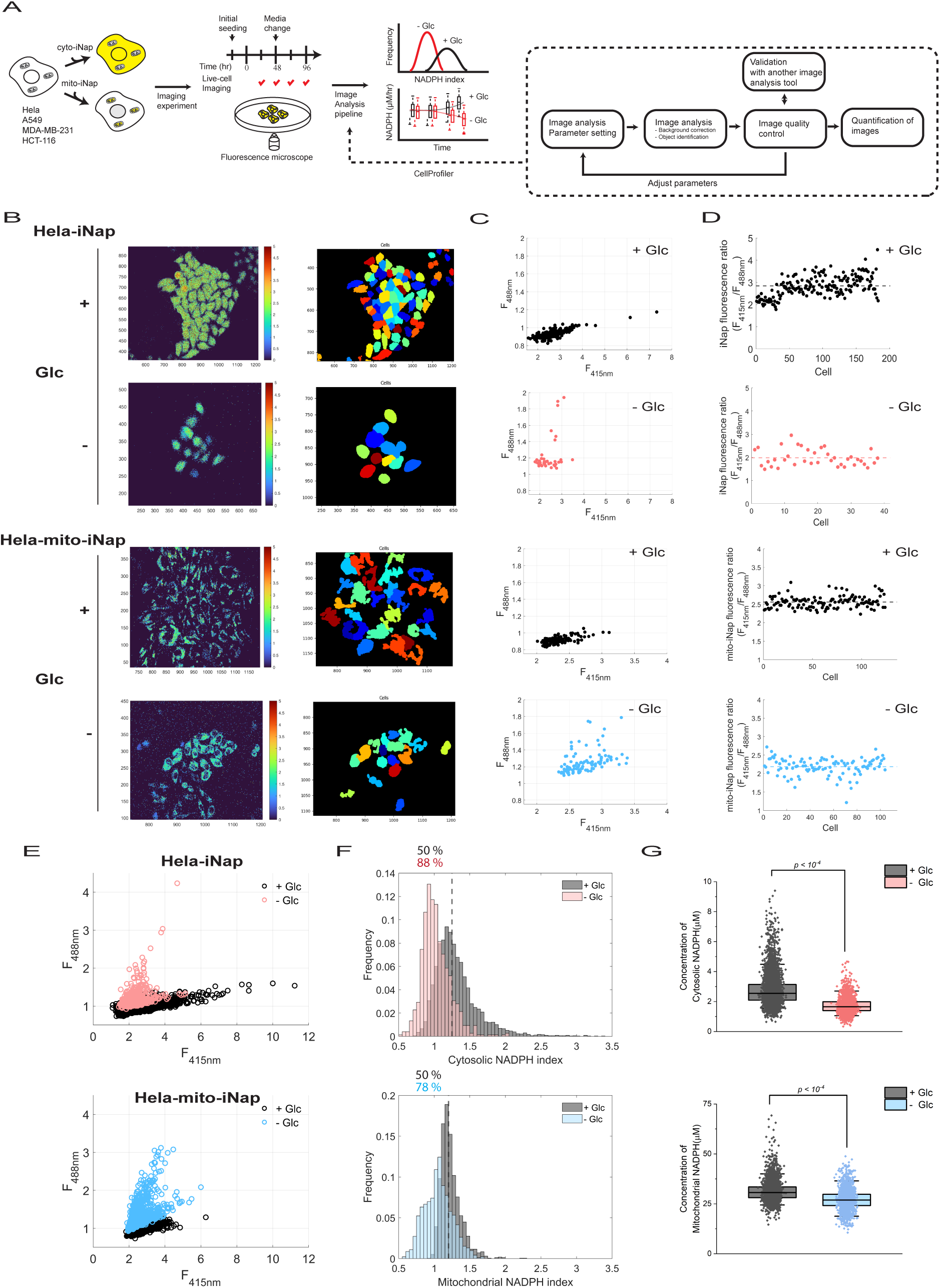
NADPH composite index analysis quantifies cytosolic and mitochondrial NADPH dynamics. **(A)** Experimental design for measurement of cytosolic and mitochondrial NADPH concentrations using iNap sensors and an automated image analysis pipeline. Cancer cell lines that stably express iNap or mito-iNap sensors are generated, and fluorescence images are obtained across multiple time points using fluorescence microscope. Subsequently, an automated image analysis pipeline that utilizes CellProfiler^29^ with optimized parameters is applied for quantification of NADPH concentration and dynamics over thousands of single cell images. **(B)** Images showing a group of Hela cells that express either iNap or mito-iNap sensors in presence or absence of glucose conditions (first columns), and identification of the cells through our image analysis pipeline (second columns). Each pixel is equal to 0.65 µm. **(C)** Scatter plots representing the quantified fluorescence intensities (ex: 415 and 488 nm) emitted from Hela-iNap or Hela-mito-iNap sensors of each identified cell in panel (B). Each dot represents single cells. **(D)** The corresponding iNap fluorescence ratios (F_415nm_/F_488nm_) in presence and absence of glucose. **(E)** Scatter plots showing the quantified fluorescence intensities (ex: 415 and 488 nm) emitted from Hela-iNap or Hela-mito-iNap sensors of all biological replicates in response to glucose deprivation (n = 3916, 1561, 917 cells for iNap+Glc, iNap-Glc, mito-iNap-Glc, mito-iNap-Glc conditions. A total three biological replicates, each of which comprising six technical replicates were used for analysis). **(F)** Histogram representing cytosolic and mitochondrial NADPH indices that are quantified in presence or absence of glucose conditions. Dashed line represents a median of the index of the control (+Glc) and the numbers above the histogram represents the percentages of the cells in each condition with the index values lower than the median of the control. **(G)** Concentration of cytosolic and mitochondrial NADPH in presence or absence of glucose. Boxplot represents the 25^th^, 50^th^, and 75^th^ percentiles with whiskers indicating 5^th^ and 95^th^ percentile. Kruskal-Wallis test was used for statistical analysis. P-values < 0.05 is considered statistically significant.

To confirm the robustness of our pipeline, we further analyzed the fluorescence ratios that were manually quantified with ImageJ, a widely used image process software developed by National Institutes of Health^30^. We found a high correlation between the measurements by two different image analysis software (R^2^ = 0.95) (***SI Appendix*, Fig. 2*A***). Overall, under glucose deprived condition, we found the fluorescence ratios of iNap and mito-iNap fluorescence ratio were approximately 23 % and 22 % lower than those of control, respectively (**Fig. 2*E* and *SI Appendix*, Fig. 2*B***). Additionally, we found the fluorescence ratio of iNap-ctr and mito-iNap-ctr were approximately 4 and 10 % lower under glucose deprived condition compared to the control, indicating that the pH increased by approximately 0.05 in cytosol and 0.1 in mitochondria based on the previously obtained pH calibration data^11^ (***SI Appendix*, Fig. 2*C*)**.

Moreover, when we compared the NADPH indices, the pH-corrected iNap sensor’s fluorescence ratio, we found that both cytosolic and mitochondrial NADPH indices were approximately 21 and 12 % lower than those of control **(Fig. 2*F*)**. We further converted the cytosolic and mitochondrial NADPH indices to the concentration by calibrating the sensors as previously described (***SI Appendix*, Fig. 2*D***)^11,12^. The conversion indicated the median concentration of cytosolic NADPH decreased by approximately 36 % from 2.6 ± 0.01 to 1.6±0.01 µM, while the mitochondrial NADPH decreased by approximately 12 % from 30.7±0.09 to 26.9±0.12 µM (**Fig. 2*G***). Altogether, the integration of iNap sensors and an image analysis pipeline demonstrated its robustness for quantifying a large number of single-cell images obtained from iNap sensors, and its application of measuring cytosolic and mitochondrial NADPH dynamics in response to nutrients perturbation.

### Lowering extracellular glucose generally decreases both cytosolic and mitochondrial NADPH levels

Next, using this pipeline, we evaluated how nutrients availability influenced cytosolic and mitochondrial NADPH dynamics. Among various nutrients, we focused on glucose as it serves as a primary fuel source for many cancer cells^31^. Glucose can regenerate cytosolic NADPH through the pentose phosphate pathway, ME1, or IDH1, and mitochondrial NADPH through ME2, IDH2, or nicotinamide transhydrogenase (NNT) (**Fig. 3*A*)**. To determine the impact of glucose availability on compartmentalized NADPH levels, we measured the cytosolic and mitochondrial NADPH indices of Hela cells cultured in DMEM with varying glucose concentrations (25, 6, 1, and 0 mM). DMEM with 25 mM glucose was used as a control. In response to varying glucose conditions, we observed different dynamics of cytosolic and mitochondrial NADPH indices. Specifically, after 48 hours, we observed that both cytosolic and mitochondrial NADPH indices started to decline in response to lower glucose concentration in media, and further exacerbated after 72 hours (**Fig. 3*B*)**. We also analyzed the spent media and found the glucose was nearly depleted in the 1 mM glucose condition after 96 hours, leading to similar responses as the 0 mM glucose condition (***SI Appendix*, Fig. 3*A*)**.

**Figure 3.**
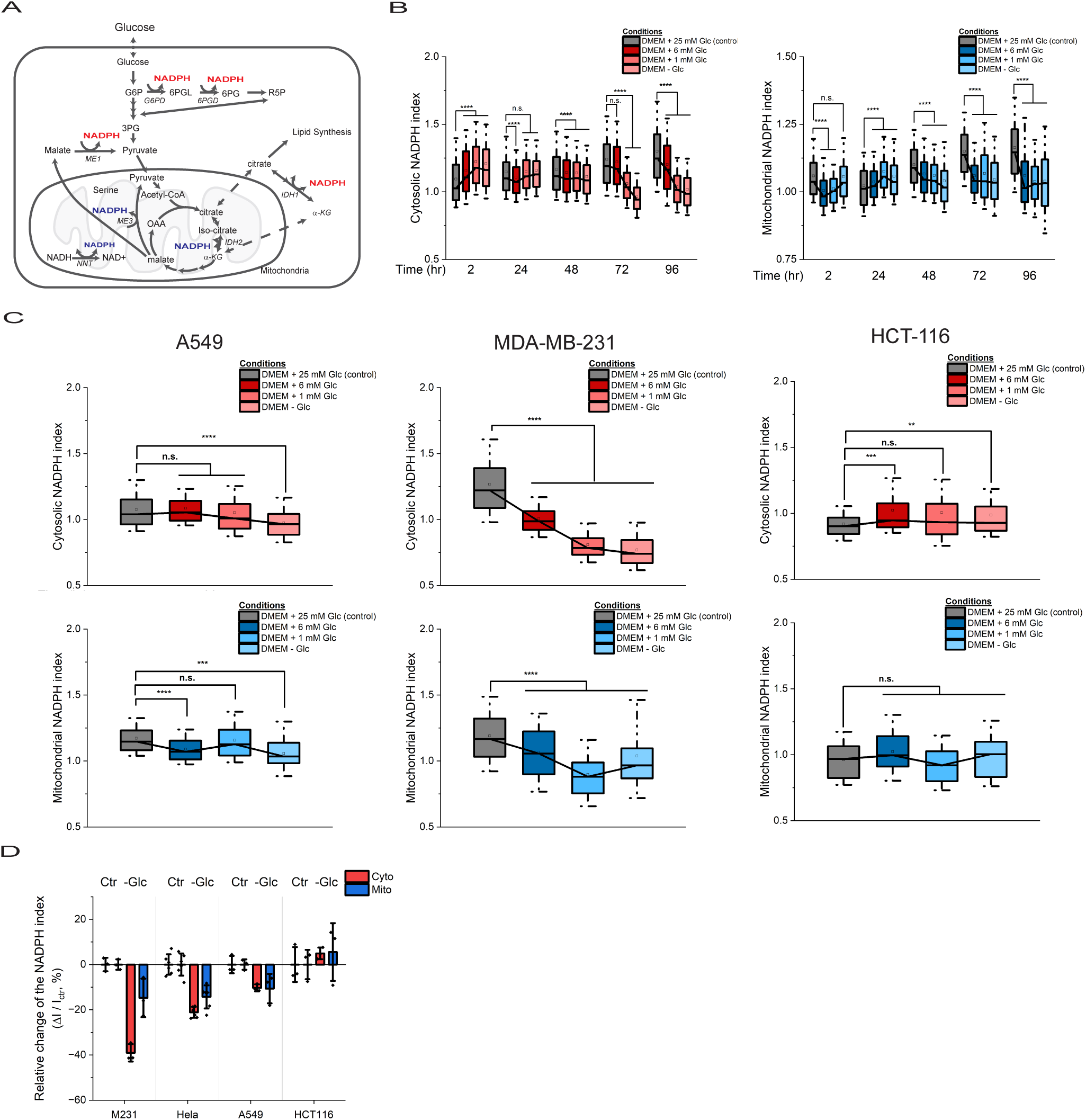
Lowering extracellular glucose generally decreases both cytosolic and mitochondrial NADPH levels. **(A)** Schematics representing glucose metabolism and enzymatic reactions that generate NADPH in cytosol and mitochondria. **(B)** Cytosolic (red) and mitochondrial (blue) NADPH indices in Hela cells cultured in varying glucose concentrations in media over 96 hours. Boxes represent the 25^th^, 50^th^, 75^th^ percentiles and the whiskers represents for 5^th^ and 95^th^ percentile. Total number of cells analyzed were found in the extended data figure 3.**(C)** Cytosolic and mitochondrial NADPH indices in A549, MDA-MB-231, and HCT-116 under varying glucose concentrations in media after 96 hours. **(D)** Relative change of cytosolic and mitochondrial NADPH indices in absence of glucose across four different cancer cell lines. Each dot represents the biological replicate.

Moreover, to evaluate whether the responses of cytosolic and mitochondrial NADPH dynamics were similar across other cancer cells of different lineages, we stably expressed iNap and mito-iNap sensors and measured compartmentalized NADPH indices in cancer cell lines, including A549 (non-small cell lung cancer), HCT-116 (colorectal cancer), and MDA-MB-231 (breast cancer) **(*SI Appendix*, Fig. 3*B*)**. Across these cell lines, we generally observed a decrease in both cytosolic and mitochondrial NADPH indices in response to reduced glucose levels (**Fig. 3*C*)**. Notably, for A549, the cytosolic and mitochondrial NADPH indices were approximately 7 % and 10 % lower than those of the control in the glucose-depleted condition. For MDA-MB-231, we found a more pronounced effect as the cytosolic and mitochondrial NADPH indices were approximately 40 % and 17 % lower, respectively. However, for HCT-116, we found the cytosolic and mitochondrial NADPH indices were around 8 % and 4 % higher (**Fig. 3*D*)**. These results indicate the absence of lower amount of glucose in media generally decreased both cytosolic and mitochondrial NADPH levels across cancer cell lines.

### Serine and glutamine deprivation influences compartmentalized NADPH dynamics differentially across cancer cell lines

Next, given that extracellular serine can be utilized to generate cytosolic or mitochondrial NADPH via the folate-dependent one-carbon metabolism pathway^5,14,32^, we assessed its impact on compartmentalized NADPH pools **(Fig. 4*A*)**. For Hela cells, we found that both cytosolic and mitochondrial NADPH indices were approximately 5 % and 7 % lower than those of the control after 96 hours of serine deprivation **(Fig. 4*B*)**. For A549, we found cytosolic NADPH index was approximately 7 % lower than that of control, while the mitochondrial NADPH index was unchanged. For MDA-MB-231, the cytosolic and mitochondrial NADPH indices were approximately 30 % and 7 % lower than those of control. Interestingly, compared to glucose deprivation, the change of cytosolic NADPH index of MDA-MB-231 cells were more pronounced than the change of mitochondrial NADPH index. For HCT-116, we found the cytosolic and mitochondrial NADPH indices were approximately 25 % and 11 % higher than those of control **(Fig. 4*C*)**. These data indicated that the impact of serine deprivation was generally not as strong as that of glucose deprivation, and the extent of the change of cytosolic NADPH levels was greater than that of mitochondrial NADPH levels.

**Figure 4.**
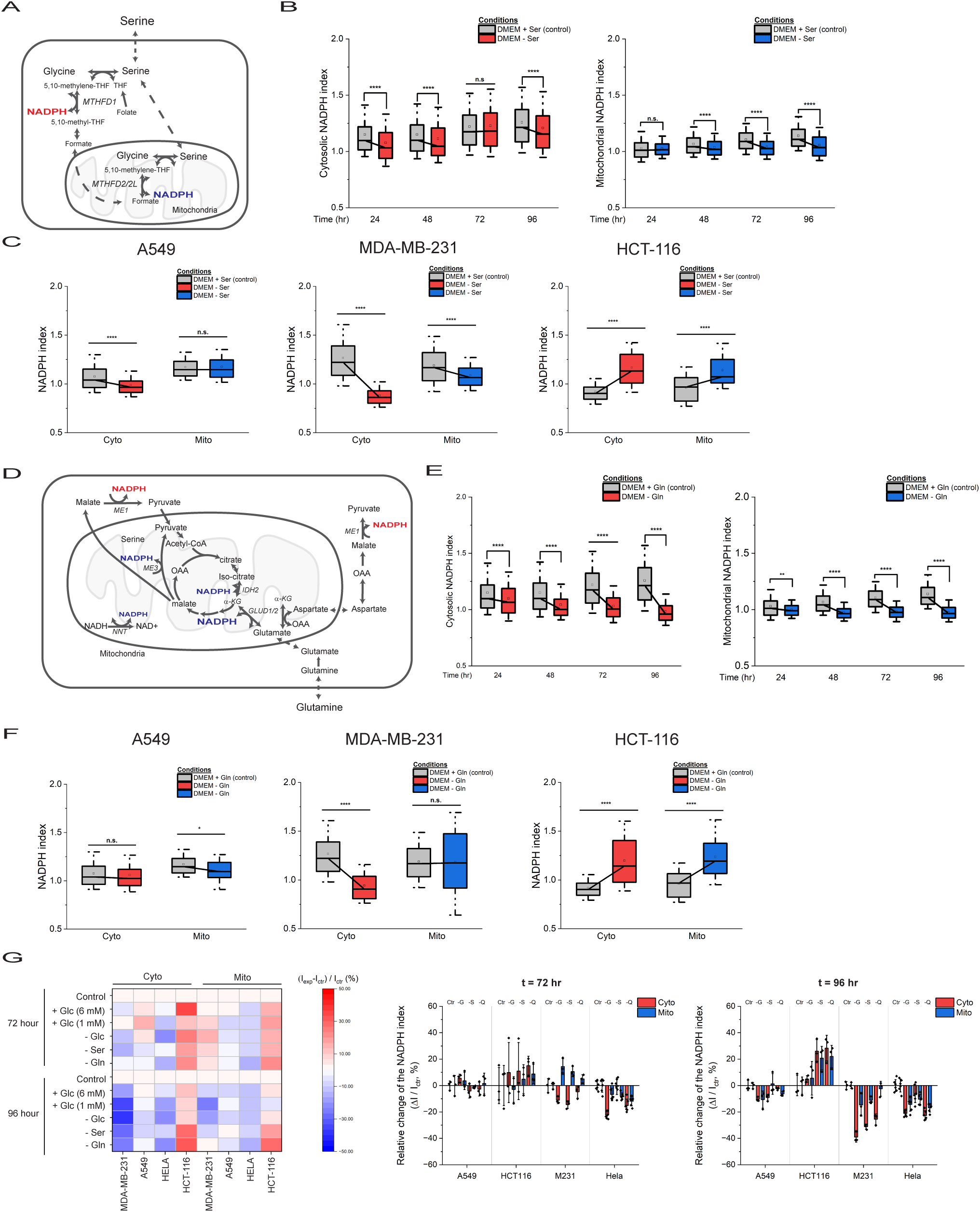
Serine and glutamine deprivation influences compartmentalized NADPH dynamics differentially across cancer cell lines. **(A)** Schematic representing the folate dependent serine metabolism that generate NADPH in cytosol and mitochondria. **(B)** Cytosolic (red) and mitochondrial (blue) NADPH indices in Hela cells cultured in absence of serine over 96 hours. **(C)** Cytosolic and mitochondrial NADPH indices in A549, MDA-MB-231, and HCT-116 in absence of serine after 96 hours. **(D)** Schematic representing the glutamine metabolism that generate NADPH in cytosol and mitochondria. **(E)** Cytosolic (red) and mitochondrial (blue) NADPH indices in Hela cells cultured in absence of glutamine over 96 hours. **(F)** Cytosolic and mitochondrial NADPH indices in A549, MDA-MB-231, and HCT-116 in absence of glutamine after 96 hours. **(G)** Heatmap and bar graphs representing a relative change of the median of cytosolic or mitochondrial NADPH indices against the index of control at 72 and 96 hours. The control condition represents cells cultured in DEME with 25 mM glucose.

Moreover, as glutamine can also regenerate NADPH via the glutaminase reaction in mitochondria or transaminase reactions followed by the malic enzyme in the cytosol^18,33^, we further evaluated the impact of glutamine deprivation on compartmentalized NADPH dynamics **(Fig. 4*D*)**. For Hela, we found that both cytosolic and mitochondrial NADPH indices were approximately 22 % and 13 % lower than those of control in absence of glutamine **(Fig. 4*E*)**. For A549, we observed minimal changes for both cytosolic and mitochondrial NADPH indices. Interestingly, for MDA-MB-231, we found that the cytosolic NADPH index was approximately 26 % lower than that of control, while the mitochondrial NADPH index was minimally perturbed. For HCT-116, we found both cytosolic and mitochondrial NADPH indices were nearly 27 % and 23 % higher than those of control, respectively **(Fig. 4*F*)**. These results suggested that the influence of glutamine on cytosolic and mitochondrial NADPH levels were dependent of cancer cell lines. Altogether, our investigations on nutrients availability revealed distinct compartmentalized NADPH dynamics across different cancer cell lines in response to glucose, serine or glutamine deprivation **(Fig. 4*G* and *SI Appendix*, Fig. 4)**.

### Cytosolic and mitochondrial NADPH dynamics patterns are cell-type dependent

Next, to estimate the rate of change of NADPH under varying nutrient conditions, we compared cytosolic and mitochondrial NADPH indices measured at 48 and 96 hours. In the absence of glucose, we found that the rate of change of cytosolic and mitochondrial indices decreased more significantly than other conditions (***SI Appendix*, Fig. 5*A***). The withdrawal of serine or glutamine also resulted in a reduction of the rate of change of NADPH indices over time, with HCT-116 being an exception. When we plotted the relative change of cytosolic and mitochondrial NADPH indices, we found distinct compartmentalized NADPH dynamics patterns in response to nutrient variations **(Fig. 5*A*)**.

**Figure 5.**
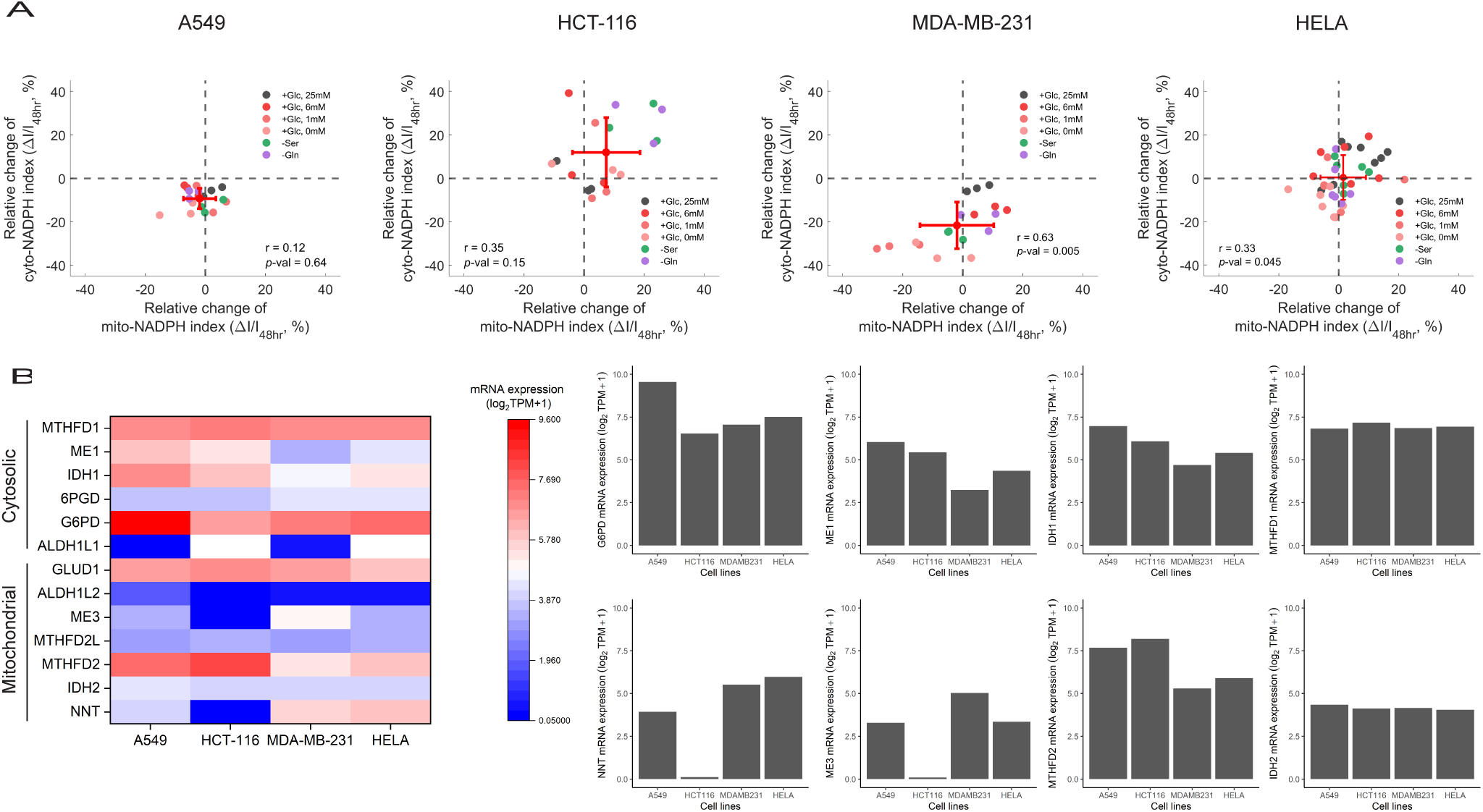
Cytosolic and mitochondrial NADPH dynamics are cell-type dependent. **(A)** Characterization of the relationship between cytosolic and mitochondrial NADPH indices under different nutrient conditions. The ΔI represents the difference between NADPH index measured at 48 and 96 hours. Each red dot with error bars represents the mean and standard deviations of all data points, each of which representing the mean of at least three biological replicates under indicated conditions. Pearson correlation coefficient and p-values were calculated. **(B)** Gene expression levels of cytosolic and mitochondrial NADPH generating enzymes were extracted from the Cancer Cell Line Encyclopedia (CCLE). Bar chart represents individual gene expression levels of *G6PD, ME1, IDH1, MTHFD1, NNT, ME3, MTHFD2,* and *IDH2* across A549, HCT-116, MDA-MB-231, and Hela.

To understand what contributed to the differential NADPH responses across cancer cell lines, we analyzed gene expression patterns of NADPH-producing enzymes from the Cancer Cell Line Encyclopedia (CCLE) (**Fig. 5*B* and *SI Appendix*, Fig. 5*B*)**^34^. We found that A549 exhibited higher *G6PD* expression compared to other cell lines, whereas HCT-116 exhibited minimal *NNT* expression. The expression levels of other genes such as *MTHFD1* and *IDH2* were relatively consistent across the cell lines. In particular, the expression levels of *NNT* were positively correlated with either cytosolic or mitochondrial NADPH indices among cancer cell lines, suggesting that NNT gene expression may reflect the baseline compartmentalized NADPH levels of individual cells. On the other hand, we found the expression levels of *MTHFD1* were negatively correlated with mitochondrial NADPH index, suggesting its level may reflect the baseline mitochondrial NADPH level of individual cell lines **(*SI Appendix*, Fig. 5*C* and *D***). Altogether, these results demonstrated the distinct compartmentalized NADPH dynamics profiles across the tested cancer cell lines, which were linked to variation of gene expression levels of NADPH-producing enzymes such as *MTHFD1* and *NNT*.

### Fractional contribution parameter serves as a guide to target cytosolic or mitochondrial NADPH metabolism for selective cancer cell growth inhibitions

Next, to quantify the relationship between the cytosolic and mitochondrial NADPH dynamics with growth rates, we introduced the NADPH composite index and a fractional contribution parameter (F) **(Fig. 6*A*)**. F can be any number between 0 and 1. Given that highly proliferating cancer cells increase NADPH production and maintain high level of cellular NADPH^35–37^, we solved the F value by maximizing the correlation between the NADPH composite index and growth rates. Indeed, we measured the concentration of cellular NADPH levels under varying nutrient conditions and found a high correlation between the NADPH level and growth rate (r = 0.93), confirming our assumption **(*SI Appendix*, Fig. 6*A*)**. For Hela cells, we determined the optimal F value (F*) as 0.57, suggesting that the correlation between composite NADPH index and growth rate was contributed by 57 % from cytosolic NADPH index and 43 % from mitochondrial NADPH index **(Fig. 6*B* and *SI Appendix*, Fig. 6*B-D*)**.

**Figure 6.**
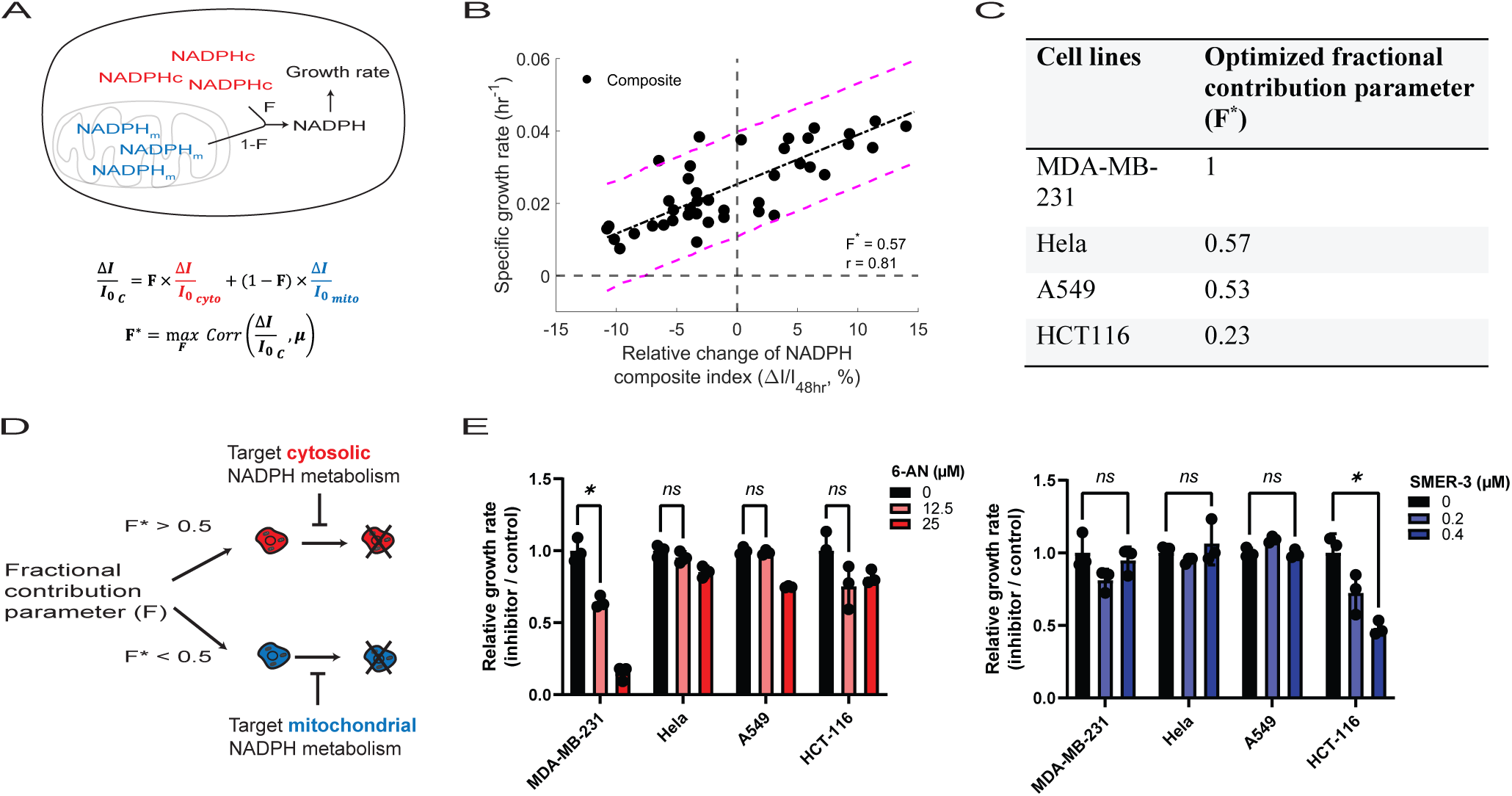
Fractional contribution parameter serves as a guide to target cytosolic or mitochondrial NADPH metabolism for selective cancer cell growth inhibitions. **(A)** Schematics describing a relationship between NADPH composite index and growth rate. The relative change of NADPH composite index (ΔI/I_0_,_C_) is defined as a linear combination of the relative change of cytosolic and mitochondrial NADPH indices (ΔI/I_0_,_cyto_ and ΔI/I_0_,_mito_, respectively) with a fractional contribution parameter (F). The optimal F is determined by maximizing the correlation between the relative change of NADPH composite index (ΔI/I_0_,_C_) and growth rates (µ). **(B)** Specific growth rate is shown as a function of the relative change of NADPH composite index with an optimized F value. Each circle represents a mean value of at least six technical replicates obtained from varying nutrient conditions including deprived glucose, serine, and glutamine. The Pearson correlation analysis was conducted and visualized using line plots with 95 % confidence intervals. **(C)** Summary of the optimized fractional contribution parameter (F*) across tested cancer cell lines. **(D)** Schematic describing the application of the fractional contribution parameter as a guide to target either cytosolic or mitochondrial NADPH metabolism. F* > 0.5 suggests to target cytosolic NADPH metabolism, whereas F* < 0.5 indicates to target the mitochondrial NADPH metabolism. **(E)** Relative growth rates of different cancer cell lines upon treatment of 6-AN, a pentose phosphate pathway inhibitor in cytosol, or SMER-3, a small molecule that generates mitochondrial oxidative stress that cause decrease in mitochondrial NADPH levels. Statistical significance was determined using multiple unpaired t-tests and p-values are provided, with a threshold of <0.05 considered statistically significant and indicated by asterisks (*P<0.05).

To test whether different cancer cell lines exhibit distinct F*, we calculated F* values across cancer cell lines. Indeed, we found their differential relationship between compartmentalized NADPH dynamics and growth rates. HCT-116 cells exhibited a negative correlation between the change of cytosolic and mitochondrial NADPH indices and growth rate, while A549 revealed minimal correlation between compartmentalized NADPH dynamics and growth rate. MDB-MB-231 showed higher correlation between cytosolic NADPH index and growth rate (***SI Appendix*, Fig. 6*E*)**. Subsequently, we quantified F* and determined its value of 0.53 for A549, 0.23 for HCT-116, and 1 for MDA-MB-231 (**Fig. 6*C* and *SI Appendix*, Fig. 6*F*)**.

Based on these observations, we hypothesized that cells with a higher F* value would be more sensitive to perturbations of cytosolic NADPH generation pathways, whereas F* closer to 0 would be more sensitive to perturbations of mitochondrial NADPH generation pathway. **(Fig. 6*D*)**. To test the hypothesis, we treated cell lines with 6-AN, an inhibitor of G6PD/6PGD in the oxidative pentose phosphate pathway, to perturb cytosolic NADPH metabolism. Upon treatment of 25 µM of 6-AN, we found the growth rate of MDA-MB-231, whose F value was 1, decreased by nearly 85 % while that of other cell lines reduced by less than 25 %. On the other hand, when we treated cells with 0.4 µM of SMER-3, a small molecule known to produce oxidative stress in mitochondria and decrease mitochondrial NADPH pool^38^, we found the growth rate of HCT-116, whose F value was 0.23, decreased by nearly 52 % while that of other cell lines reduced by less than 5 % (**Fig. 6*E*)**. Altogether, these findings demonstrated the utility of the fractional contribution parameter in quantifying cell-type specific growth responses to interventions that target compartmentalized NADPH metabolism.

### Citrate transporter, encoded by *SLC25A1*, is essential for maintaining compartmentalized NADPH pools and cell growth

As the NADPH composite index reflected the contribution of both cytosolic and mitochondrial NADPH pools, we were wondering which mitochondrial transporters played key roles in shuttling the compartmentalized NADPH pools. To determine a key mitochondrial NADPH transporter for NADPH, we performed *in-silico* gene-knockout screening on *SLC25* gene families using a curated genome-scale metabolic model (GEM) (**Fig. 7*A*)**. First, we reconstructed Hela-GEM by using RNA-seq data for Hela^39^ and a task-driven integrative network inference for tissue (tINIT) algorithm^40^. Next, we analyzed the spent media and calculated the extracellular uptake rates of carbon sources and amino acids, which were used as constraints on exchange rates in the model (***SI Appendix*, Fig. 7*A***). After flux balance and variability analysis on the curated model, we performed Monte Carlo random sampling analysis to determine flux distributions on knockout simulations of *SLC25* genes. The analysis revealed that the ablation of *SLC25A1* resulted in the most significant decrease in the net generation rate of NADPH compared to wildtype and other *SLC25* knockouts (**Fig. 7*B***). The net NADPH generation rate was predicted to be nearly 56 % lower than that of the wildtype (**Fig. 7*C*)**. The rate through the pentose phosphate pathway, IDH1, and MTHD1 were also predicted to decrease in *SLC25A1* knockout cells (**Fig. 7*D* and *SI Appendix*, Fig. 7*B*).**

**Figure 7.**
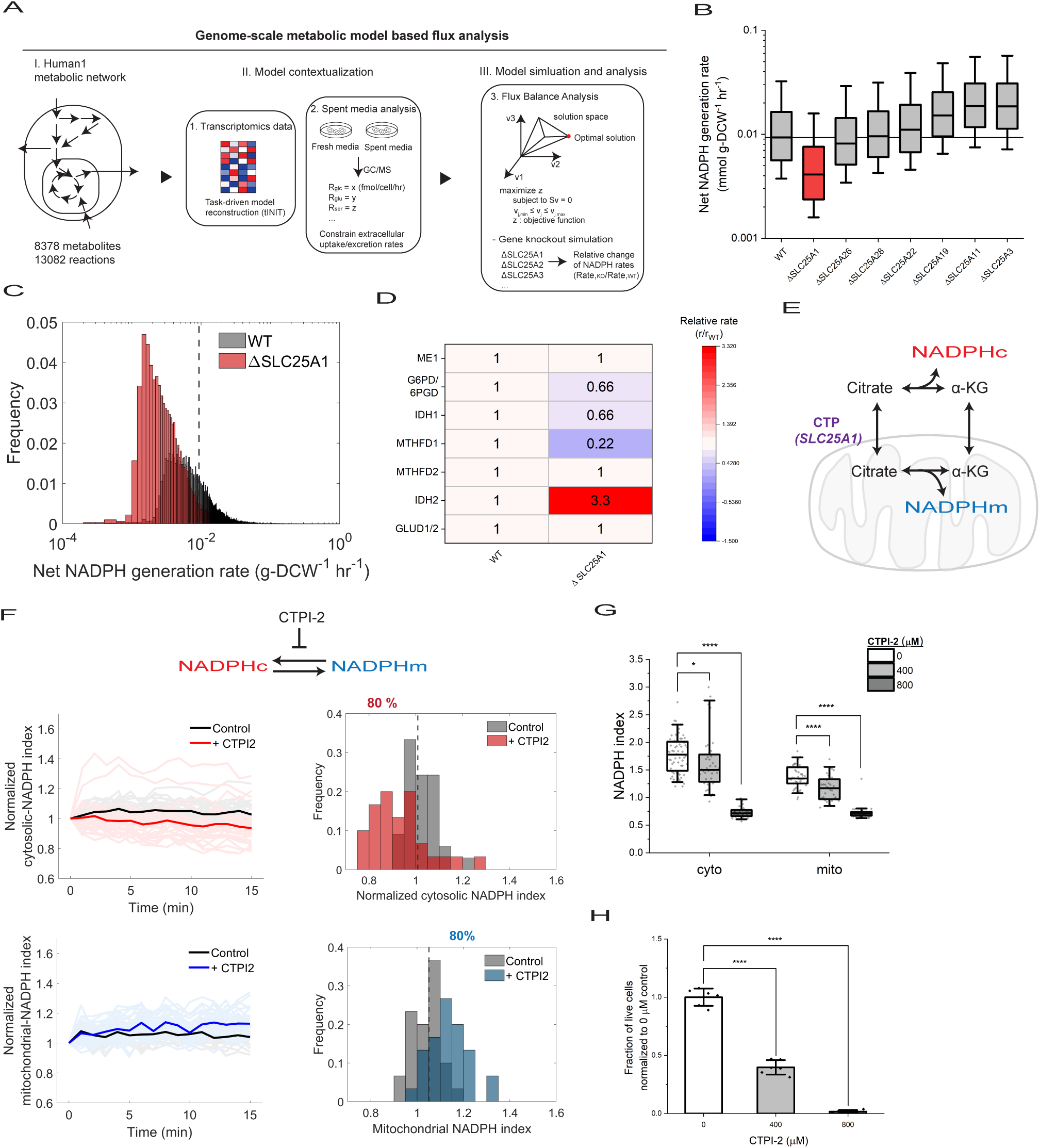
Citrate transporter, encoded by *SLC25A1*, is essential for maintaining compartmentalized NADPH pools and cell growth. **(A)** Overview of the genome-scale metabolic model (GEM) based flux and knockout analysis. Hela-specific genome-scale metabolic model was reconstructed based on the gene expression levels of Hela cells. The spent media was analyzed to calculate the uptake rates and used as the constraints to the model. Flux balance, variability, sampling analyses were performed in response to gene knockout of *SLC25* gene family. **(B)** Simulated knockout analysis on *SLC25* genes and the corresponding net NADPH generation rates. Boxplot represents the 25^th^, 50^th^, and 75^th^ percentiles with whiskers indicating 5^th^ and 95^th^ percentile (n_sampling_ = 14,848). **(C)** Distribution of the predicted net generation rates of NADPH for wild-type and *SLC25A1* knockouts. Dashed line represents a median rate of the wild-type. **(D)** Mean reaction rates of NADPH regenerating enzymes upon knockout of *SLC25A1*. **(E)** Schematic illustrating the indirect NADPH shuttle system facilitated by the citrate transporter (CTP, encoded by *SLC25A1*). **(F)** Time-course measurement of cytosolic (red) and mitochondrial (blue) NADPH indices upon inhibition of citrate transporter with 400 µM of CTPI-2 compound. Histogram represents a distribution of cytosolic and mitochondrial NADPH indices after 15 minutes of treatment of the inhibitor. Dashed line represents a median of the index of the control and the numbers indicate the percentage of cells with NADPH indices lower or higher than the median of the control (n = 30 cells for each condition in two biological replicates). **(G)** Cytosolic and mitochondrial NADPH indices were measured upon treatment of 400 and 800 µM CTPI-2 after 24 hours (n = 65, 40, 37, 43, 37, 31 in three biological replicates). **(H)** Fraction of live cells after treatment of 400 and 800 µM CTPI-2 for 24 hours (n = 6).

Functionally, *SLC25A1* encodes a citrate transporter that carries citrate and iso-citrate across mitochondrial membrane and shuttle NADPH through cycling reactions (**Fig. 7*E***). To test the model-based analysis, we measured the cytosolic and mitochondrial NADPH dynamics after treating the cells with citrate transporter inhibitor (CTPI-2). After 15 minutes of inhibition of citrate transporter, the cytosolic NADPH index was nearly 9 % lower than that of control, whereas the mitochondrial NADPH index was nearly 10 % higher (**Fig. 7*F*)**. After 24 hours of treatment of 800 µM CTPI-2 treatment, we found that both cytosolic and mitochondrial NADPH indices were nearly 60 % and 47 % lower than those of control, respectively **(Fig. 7*G*)**. Inhibition of the citrate transporter also inhibited cell growth **(Fig. 7*H*)**.

Additionally, to investigate the role of *SLC25A1* gene in NADPH metabolism and growth rates, we performed gene essentiality, co-essentiality, and gene enrichment analysis by using the genome-scale CRISPR knockout screening data from the Broad Institute^41^ **(*SI Appendix*, Fig. 7*C*)**^41^. We found that knocking out of *SLC25A1* generally decreased cell growth rates across 739 cell lines (***SI Appendix*, Fig. 7*D* and Table 1**). Furthermore, a co-essentiality analysis and biological process enrichment analysis revealed that *SLC25A1* showed a high correlation with genes that are associated with NAD(P)^+^ oxidoreductase activities and nucleotide biosynthetic processes (***SI Appendix*, Fig. 7*E* and Table 2)**. Altogether, our data suggests that citrate transporter, encoded by *SLC25A1*, plays key roles in maintaining the compartmentalized NADPH pools and cell growth, as its inhibition decreased both compartmentalized NADPH pools and growth rates.

## Discussion

Here, we present NADPH composite index analysis, which quantifies the relationship between cytosolic and mitochondrial NADPH dynamics to growth rates in response to varying nutrients. Although previous studies focused on determining pathways that were important for NADPH regeneration, less is known about how compartmentalized NADPH dynamics are directly linked to growth rates and whether these relationships could be quantified across different cancer cell lines. Our results not only demonstrated differential relationship between compartmentalized NADPH dynamics and growth rates across varying cancer cell lines, but also demonstrated its utility to selectively inhibit cancer cell growths by targeting cytosolic or mitochondrial NADPH metabolism.

Specifically, we demonstrated how nutrient availability, particularly glucose, serine, and glutamine, influenced cytosolic and mitochondrial NADPH dynamics. Although glucose’s role in supplying the cytosolic NADPH pool via the pentose phosphate pathway is well-documented^15^, its effect on mitochondrial NADPH level has remained less clear. Our findings indicated that glucose is critical for maintaining mitochondrial NADPH levels across the cancer cell lines.

Additionally, while we previously found that mitochondrial NADPH level was nearly 50 % lower in response to mitochondrial oxidative stress in Hela cells ^12,42^, we found the mitochondrial NADPH levels was around 10 % lower in the absence of glucose. This suggests that mitochondrial NADPH and its antioxidant network system may not be completely collapsed in Hela even under glucose deprived condition. This could be attributed by the compensatory mechanisms that allowed regeneration of mitochondrial NADPH through other metabolic pathways.

As serine and glutamine were reported as important sources of NADPH in cytosol and mitochondria, we further evaluated the impacts of these amino acids to compartmentalized NADPH pools. In general, we found that the absence of extracellular serine lowered both cytosolic and mitochondrial NADPH across cancer cell lines. In Hela, withdrawal of serine lowered the mitochondrial NADPH more than the cytosolic NADPH, consistent with previous studies, which indicated the serine mediated one carbon metabolism generated mitochondrial NADPH^5,43^. Additionally, we found cytosolic and mitochondrial NADPH levels in MDA-MB-231 were greatly influenced by the removal of serine. This supports the finding that MDA-MB-231 is known to rely on extracellular serine for proliferation due to its low expression of serine synthesis pathway^44^. Similarly, withdrawal of glutamine also generally decreased the cytosolic and mitochondrial NADPH pools. The decrease of mitochondrial NADPH levels could be attributed by decreased reaction rate of glutamate dehydrogenase (GLUD1), which converts glutamate to α-KG, while regenerating NADPH in mitochondria^45^. For cytosolic NADPH, the conversion of glutamate to α-KG through aspartate transaminase (GOT1 and GOT2) and the subsequent malic enzyme (ME1) reaction could generate NADPH in both cytosol and mitochondria^18^. While GLUD reaction was dispensable in presence of glucose^46^, our results demonstrated that the change of cytosolic and mitochondrial NADPH levels were dependent on cancer cell types, suggesting a need to characterized compartmentalized NADPH levels for individual cancer cells, as the mitochondrial NADPH levels of MDA-MB-231 cell lines were less sensitive to glutamine deprivation.

Furthermore, regarding how cytosolic and mitochondrial NADPH could be transported across the compartment, we identified the citrate transporter, encoded by *SLC25A1*, as a key mitochondrial transporter that supported the maintenance of compartmentalized NADPH pools. Previous studies focused on specific mitochondrial transporters for NADPH metabolism study, but our study improved this limit by comparatively and simultaneously analyzing all mitochondrial transporters through *in-silico* analysis^47^. Although *SLC25A1* was previously reported in its role in shuttling NADPH from cytosol to mitochondria through indirect metabolite shuttle systems in context of anchorage independent growth ^22^, it was unclear how the citrate transporter directly impacted the cytosolic and mitochondrial NADPH pools. Our results demonstrated that inhibition of citrate transport decreased both cytosolic and mitochondrial NADPH in the long term, suggesting the decrease in cellular NADPH/NADP^+^ ratio as observed in the previous studies were likely contributed by the decrease in both cytosolic and mitochondrial NADPH pools.

Finally, as maintaining cytosolic and mitochondrial NADPH redox states are critical for survival and proliferation^48^, we quantified the relationship between the compartmentalized NADPH dynamics and growth rates by introducing NADPH composite index and a fractional contribution parameter (F). This parameter not only determined the relative contribution of cytosolic and mitochondrial NADPH dynamics to growth rates, but also demonstrated its utility as a guide to target cytosolic or mitochondrial NADPH metabolism for cancer-specific selective growth inhibitions. Future studies may expand our NADPH composite index analysis to investigate other cancer cell types and test the effectiveness of targeting cytosolic and mitochondrial NADPH metabolism *in vivo*. In summary, we introduce NADPH composite index analysis to quantify the relationship between compartmentalized NADPH dynamics and growth rates, highlighting its potential to serve as a guide to target cytosolic or mitochondrial NADPH metabolism for selective cancer cell growth inhibitions.

## Materials and Methods

### Creation of stable cell lines expressing iNap33, iNapC, mito-iNap, and mito-iNapC sensors

HEK293 cells were seeded at 7.5 × 10^5^ cells per 35mm well in 6 well plates (Corning, VWR 29442-042) for two days until 70 % – 90 % confluency. pLJM-1 vectors encoding appropriate sensors were co-transfected with the packaging plasmids pMD2.G and Pax2 vectors at a 3: 1: 2 ratio for a total of 5 μg of sensor containing plasmids and 10 μg of Lipofectamine 2000 in OptiMEM medium overnight. Next morning, the media was replaced with 1mL of Dulbecco’s modified Eagle’s medium (DMEM; Lonza) supplemented with 10% fetal bovine serum (FBS; ATCC) and the media was collected every 24 hour for two days. The collected media was centrifuges at 500 g for five minutes and the supernatant was collected and stored at -80C freezer. After virus containing supernatant was prepared, cell lines (Hela, A549, HCT-116, and MDA-MB-231) were seeded at 7 × 10^5^ cells per 35 mm well in 6 well plates for a day until 70 % – 90 % confluency. 1 mL of virus-containing supernatant was added to wells containing each cell lines with 6 μg/mL of polybrene. After three days of infection, cells from each well were expanded to 10 cm dish with 6 μg/mL puromycin selection media for about seven days until 70-90% confluency.

### Cell culture

All cell lines were grown in Dulbecco’s modified Eagle’s medium (DMEM; Lonza) supplemented with 10% fetal bovine serum (FBS; ATCC) at 37 °C with 5% CO2. For DMEM media with varying concentration of glucose, we reconstituted the media by dissolving the DMEM powder that contained no glucose (Cat # D9800-26) and added back glucose to make a final concentration of 25 mM, 6 mM, 1 mM, and 0 mM glucose. Similarly, for DMEM media without serine or glutamine, we reconstituted the media and added back serine or glutamine to make a final concentration of 0.4 mM and 4 mM, respectively. The reconstituted media were sterile filtered with a vacuum filtration system with membranes of 0.20 µm pore size.

### Determination of specific growth rates

Cells were plated at 20,000 (Hela, A549, HCT-116) or 40,000 cells per well in 6-well plates and cultured in 2 mL DMEM with 10 % FBS overnight. After a day, one well was used to count the initial cell number using a Cellometer (Nexcelom Bioscience). In other wells, cells were washed twice with 2 mL PBS and 2 mL of fresh media for experimental conditions was added. After two days, the media replaced to a fresh 2 mL of media to prevent depletion of media components. After four days, final cell number was again counted using a Cellometer. Specific growth rates (µ) were determined using the following equation:

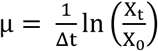

where X_t_ and X_0_ are the cell numbers s at time 0 and time t.

### Cellular imaging with fluorescence microscopy

The fluorescence emission signal at 515 nm was recorded using an inverted IX81 widefield fluorescence microscope (Olympus) with a 20x objective lens and Prior Lumen 2000 lamp. The Chroma 415/30 nm and a Semrock 488/6 nm excitation filters were used and a Semrock 525/ 40 nm filter was used for the emission filter. The exposure time was 300 ms and 10% lamp intensity was used. In-built multi-dimensional setup was used for automated time-course imaging recording. Images were captured every minute and exported to CellProfiler for post image processing.

### Image analysis with CellProfiler

A set of raw images, which were obtained from imaging experiments with 415 and 488 nm excitations, was imported to CellProfiler image analysis software^29^. Enhance module with speckles feature was applied to enhance the intensity of pixels of the cells relative to the background noise, substituting an illumination correction step for background noise correction. Afterwards, three classes Otsu threshold algorithm was used to identify cells based on intensity thresholding. IdentifyPrimaryObjects module with a defined objects dimeter size in pixel units was used to assign the threshold-applied cells as individual objects. The average pixels values of individual cells were calculated with a MeasureImageIntensity module, followed by a calculation of the ratio between pixel values of the 415 and 488 nm. Image quality control was performed before processing all the acquired images. Parameters such as threshold smoothing scale, correction factor parameters, and diameter of objects parameter were adjusted to ensure identified objects matched to the cells showing fluorescence. Images created using the pipeline and the extracted features were exported to MATLAB 2020a for data analysis and visualization with pseudo colors.

### Image analysis with using ImageJ

Backgrounds of short (415nm) and long (488nm) wavelength images were subtracted using a rolling ball algorithm from ImageJ. Long-wavelength images were converted to 32 bit and a threshold of one was applied to minimize artifact. The pixels values of the 415 nm filters were divided those of the 488 nm. Individual cells, neither too bright not too dim, were randomly selected and the mean fluorescence intensity of region of interest was calculated. All the fluorescence emission ratios were recorded as the mean ± S.E.M. The images were created using the image processing algorithm in MATLAB 2016a with pseudo colors

### Determination of extracellular fluxes

Cells were seeded at 40,000 (Hela, A549, HCT-116) or 80,000 (MDA-MB-231) per well of 6-well plates and cultured in a DMEM with 10 % FBS. The following day, the media was replaced to the fresh media of experimental conditions. 1 mL of media was immediately collected, centrifuged at 500 g for 3 minutes to remove any cell debris and frozen at 80 °C for later analysis. Cell number was also counted using a Cellometer (Nexcelom Bioscience). After two days, the media was collected and the cell number was determined. Amino acids concentration was determined by GC-MS as previously described^49,50^. In brief, 10 µL of spent media samples were mixed with 10 µL of known concentration of isotopically labeled internal standards of amino acids (Cambridge Isotope Laboratory, Andover, MA, MSK-A2-1.2). 600 µL of ice cold HPLC grade methanol was added, vortexed for 10 minutes, and centrifuge at 21,000 g for 10 minutes. 450 µL of each extract was removed and dried under nitrogen gas and stored at -80C until further analysis with GC-MS. In parallel, to make standard curves, isotopically labeled internal standards of amino acids were serially diluted and prepared for GC-MS analysis. After the concentrations of amino acids were calculated using the ratios of unlabeled to labeled amino acids through the GC-MS analysis and correction for natural abundance, the extracellular uptake/release rates of amino acids were determined based on the following equation^51^:

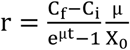

where r is the extracellular rate of amino acid, C_f_ and C_i_ are the final and initial concentration, µ is the specific growth rate, and X_0_ is the initial cell number. For concentration of glucose, lactate, glutamine, and glutamate, YSI-2900D Biochemistry Analyzer (Yellow Springs Instruments, Yellow Springs, OH) was used according to the manufacturer’s instructions and the rates were calculated.

### Gas chromatography-mass spectrometry (GC-MS) analysis for polar metabolites

GC-MS analysis for polar metabolites was described previously ^5,12^.Dried polar metabolites were derivatized by addition of Methoxyamine in pyridine (MOX) at 40 °C for 1.5 hour and *tert-*butyldimethylchlorosilane (TBDMS) was subsequently added and dried at 60 °C for 1 hour. The derivatized samples were injected on the GC/MS. GC/MS analysis was complete using an Agilent 6890 GC connected with a 30-m DB-35MS capillary column with an Agilent 5975B MS operation under electron impact ionization at 70 eV. 1 μL of the sample was injected at 270 °C using helium as the carrier gas at a flow rate of 1 mL per minute. The GC oven temperature was fixed at 100 °C for 3 minute and increased to 300 °C at 3.5 per minute. The detector was set in scanning mode and the mass-to-charge ratio was measured in range of 100 – 1000 m/z. Mass isotopomer distributions were calculated by integrating respective ion fragments of each metabolites using the in-house MATLAB code that corrected for natural abundance^52^.

### Essentiality, co-essentiality, and PANTHER pathway-enrichment analysis

The CRISPR-Cas9 knockout-out screens dataset (Achilles_gene_effect.csv) on 18333 genes in 689 cell lines was retrieved from the DepMap database (www.depmap.org). Gene scores for each SLC25 across the cell lines were extracted and visualized by a boxplot that represented median, 25^th^, and 75^th^ quartile. Co-essentiality was performed by correlating each of SLC25A gene with the rest of other genes across the cell lines and calculated a Pearson correlation coefficient. Top 50 genes with highest coefficients were exported for pathway-enrichment analysis using the PANTHER (www.pantherdb.org). Adjusted P value, reflecting the FDR, was calculated by Benjamini-Hochberg procedure and a critical value of 0.05 is used to filter results^53^.

### Genome-scale metabolic model (GEM)

A human GEM lineage (Human-GEM) was used as a core metabolic network^54^. The original model contained 13,078 reactions, 8,370 metabolites, and 3,625 genes. We incorporated RNA-seq data of Hela cells^39^ to generate Hela-specific GEM using a task-driven integrative network inference for tissue (tINIT) algorithm^40^. The contextualized Hela-GEM contained 7,424 reactions, 5,378 metabolites, and 2,251 genes.

### Flux analysis

To represent the availability of media components that impact metabolic fluxes, we experimentally obtained and constrained uptake rates of a total of 17 media components in DMEM medium, including glucose, glutamine, arginine, tyrosine, leucine, isoleucine, valine, threonine, glutamate, phenylalanine, serine, methionine, histidine, glycine, and proline, using Yellow Spring Instruments (YSI) 2950, Agilent 7890B GC and 5977 MS instrument^55^. Upon imposing constraints on the exchange rates, the flux balance analysis was implemented by maximizing the flux through biomass reaction. Given cells require high demand of NADPH supply for lipid synthesis, we fixed a half of optimized growth rate as a maximum value and subsequently performed a second optimization by maximizing the flux to NADPH production in cytosol and mitochondria.

### Gene-knockout simulation and random sampling

Single-gene knockout simulation was implemented by deleting the gene of interest. The random sampling of the solution space was determined using GpSampler algorithm, which implemented the Artificial Centering Hit-and-Run algorithm, and only those solutions that achieved fifty percent of the maximum objective function were kept^56^. A total of 14,848 random sample points was used as initial input points in sampling.

### Cellular NADPH, NADP^+^, and NADPH/NADP^+^ measurement

The measurements were performed using the NADPH/NADP^+^ Glo Assay (Promega) with a modification in manufacturer instruction. Approximately 5 x 10^5^ cells were seeded per each of well in 6-well dish in 2 mL DMEM, supplemented with 10% dialyzed FBS. After a day, the medium was aspirated and placed on ice. The cells were washed twice with 2 mL of ice-cold PBS. Then, an ice-cold 1:1 mixture of PBS and 1% dodecyltrimethylammonium bromide in 0.2 M NaOH was added to the wells, and cells were collected by scraping for 20 seconds and transferred to the ice-cold 1.5 mL centrifuge tube. The cell lysates were split to two with an equal volume of 200 μL. One aliquot was left untreated, while the other aliquot was treated with 100 μL of 0.4 M HCl. Both aliquots were placed in an incubator with 60 °C for 20 minutes. Afterwards, the manufacturer instructions were used to measure NADPH, NADP^+^, and NADPH/NADP^+^. The final concentrations of NADPH and NADP^+^ were calculated after generation of a calibration curves with known amount of NADPH and NADP^+^.

### Fractional contribution parameter

NADPH composite index is defined as I_composite_ = F × I_cyto_ + (1-F) × I_mito_, where I_composite_ represented NADPH composite index, I_cyto_ and I_mito_ represented cytosolic and mitochondrial NADPH indices, respectively. F represented a fractional contribution parameter. The equation was converted in terms of relative change and represented as 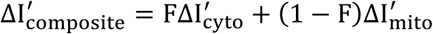, where ΔI^′^ represented the relative change of respective index from 48 hour to 96 hours. Assuming the availability of cellular NADPH is proportional to the cell growth, the specific growth rate can be expressed as function of 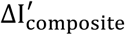. The fractional contribution parameter was calculated by maximizing the correlation between the specific growth rate and the change of NADPH index.

### Statistics and reproducibility

All statistical analysis was performed in Origin2020b, MATLAB2020b, or Prism. Number of independent technical and biological replicates, error bars and P values or statistical significances were specified in each figure legend. Technical replicates in imaging analysis represented samples obtained from different field of views in a biological replicate. Imaging experiments with box plots represented at least three biological replicates and combined into a single data set. For statistical significance analysis, Kruskal-Wallis test with Dunn-Sidak multiple comparison tests was used with an assumption that the normality was not maintained. For growth rate data, an unpaired two-tailed Student’s t-test was used. For growth inhibition experiments with small molecules, the multiple unpaired t-tests were used and p-values were provided, with a threshold of <0.05 considered statistically significant and indicated by asterisks (∗ P < 0.05, ∗∗ P < 0.01, ∗∗∗ P < 0.001,∗∗∗∗ P < 0.0001).

### Data Availability

Materials and codes can be made available from the corresponding author upon request.

## Author Contributions

Conceptualization, S.J.M., G.N.S., and H.D.S.; Methodologies, S.J.M., W.D. A.C.P.; Investigation: S.J.M., W.D., A.C.P.; Writing – Original Draft, S.J.M.; Writing – Review and Editing, S.J.M, A.C.P, W.D., J.K.K., M.G.V., G.N.S., H.D.S.; Supervision, J.K.K., M.G.V., G.N.S., H.D.S.; Funding Acquisition, S.J.M., G.N.S., H.D.S.

## Acknowledgements

We thank members of the Sikes and Stephanopoulos laboratories for useful discussions and experimental advice. We additionally thank Iva Gramatikov from Vander Heiden laboratory for assisting YSI-2900 analyzer based experiments. We also thank Yoseb Dong from Prather laboratory for valuable advice regarding genome-scale metabolic modeling. S.J.M. acknowledges support from John Henry Grover Fund in Chemical Engineering and Samsung Fellowship. G.N.S. acknowledges support from NIH R01 CA160458. H.D.S. acknowledges support from the Esther and Harold E. Edgerton endowed chair and the J.H. and E.V. Wade Fund.

