## Supplementary Figures for "NADPH composite index analysis quantifies the relationship between compartmentalized NADPH dynamics and growth rates in cancer cells"

### Supplementary materials

| REAGENT or RESOURCE | SOURCE | IDENTIFIER |
|---|---|---|
| **Bacterial and Virus Strains** | | |
| **Chemicals, Peptides, and Recombinant Proteins** | | |
| NADPH, Tetrasodium Salt | VWR | Cat # 80053-342 |
| METABOLOMICS AMINO ACID MIX STANDARD | Cambridge Isotope Lab | Cat # MSK-A2-1.2 |
| D-Glucose | Sigma-Aldrich | Cat # G8270-10KG |
| L-Glutamine | Sigma-Aldrich | Cat # G8540-100G |
| L-Serine | Sigma-Aldrich | Cat # S-4311 |
| tert-Butyldimethylsilyl chloride | Sigma | Cat # 190500 |
| Methoxamine (MOX) Reagent | ThermoFisher Scientific | Cat # TS-45950 |
| Citrate transport inhibitor (CTPI-2) | Enamine | Cat # EN300-00328 |
| Opti-MEM Media | ThermoFisher Scientific | Cat # 31985070 |
| DMEM/High: with 4.0 mM L-Glutamine, without Sodium Pyruvate. | VWR | Cat # 16750-112 |
| DMEM, no glucose, no glutamine, no phenol red | ThermoFisher Scientific | Cat # A1443001 |
| DMEM w/Sodium Bicarbonate w/o Amino Acids, Glucose, Pyruvic Acid, Phenol Red (Powder) | US biological Life Science | Cat # D9800-26 |
| Dulbecco's MEM (DMEM) w/o Glucose, Glutamine, Serine, Glycine, Sodium Pyruvate (Powder) | US biological Life Science | Cat # D9802-01 |
| MEM Non-Essential Amino Acids Solution | Thermo fisher | Cat # 11140050 |
| MEM Amino Acids Solution (50X) | ThermoFisher Scientific | Cat # 11130051 |
| **Critical Commercial Assays** | | |
| NADP/NADPH-Glo™ Assays | Promega | Cat # G9081 |
| **Experimental Models: Cell Lines** | | |
| Hela-iNap3, Hela-iNapC, Hela-mito-iNap33, Hela-mito-iNapC | This paper | N/A |
| A549-iNap3, A549-iNapC, A549-mito-iNap33, A549-mito-iNapC | This paper | N/A |
| HCT116-iNap3, HCT116-iNapC, HCT116-mito-iNap33, HCT116-mito-iNapC | This paper | N/A |
| MDA-MB-231-iNap3, MDA-MB-231-iNapC, MDA-MB-231-mito-iNap33, MDA-MB-231-mito-iNapC | This paper | N/A |
| **Recombinant DNA** | | |
| pLJM1-EGFP | Sancak et al Science. 2008 | Addgene #19319 |
| pMD2.G | Trono Lab | Addgene # 12259 |
| psPAX2 | Trono Lab | Addgene # 12260 |
| pLJM1-iNap vectors (iNap3, iNapC, mito-iNapp33, mito-iNapC) | Moon et al., 2020 | DOI: 10.1002/btm2.10184 |
| **Software and Algorithms** | | |
| MATLAB | MathWorks | www.mathworks.com |
| CellProfiler | CellProfiler | https://cellprofiler.org/ |
| ImageJ | Schneider et al., 2012 | www.imagej.net |
| Prism | GraphPad | www.graphpad.com |

### Experimental Model and Subject Details

Hela cells (ATCC® CCL- 2) and HEK-293 cells (ATCC® CRL-1573) lines belonged to Sikes laboratory. MDA-MB-231, A549 and HCT116 cell lines belonged to Greg Stephanopoulos laboratory.

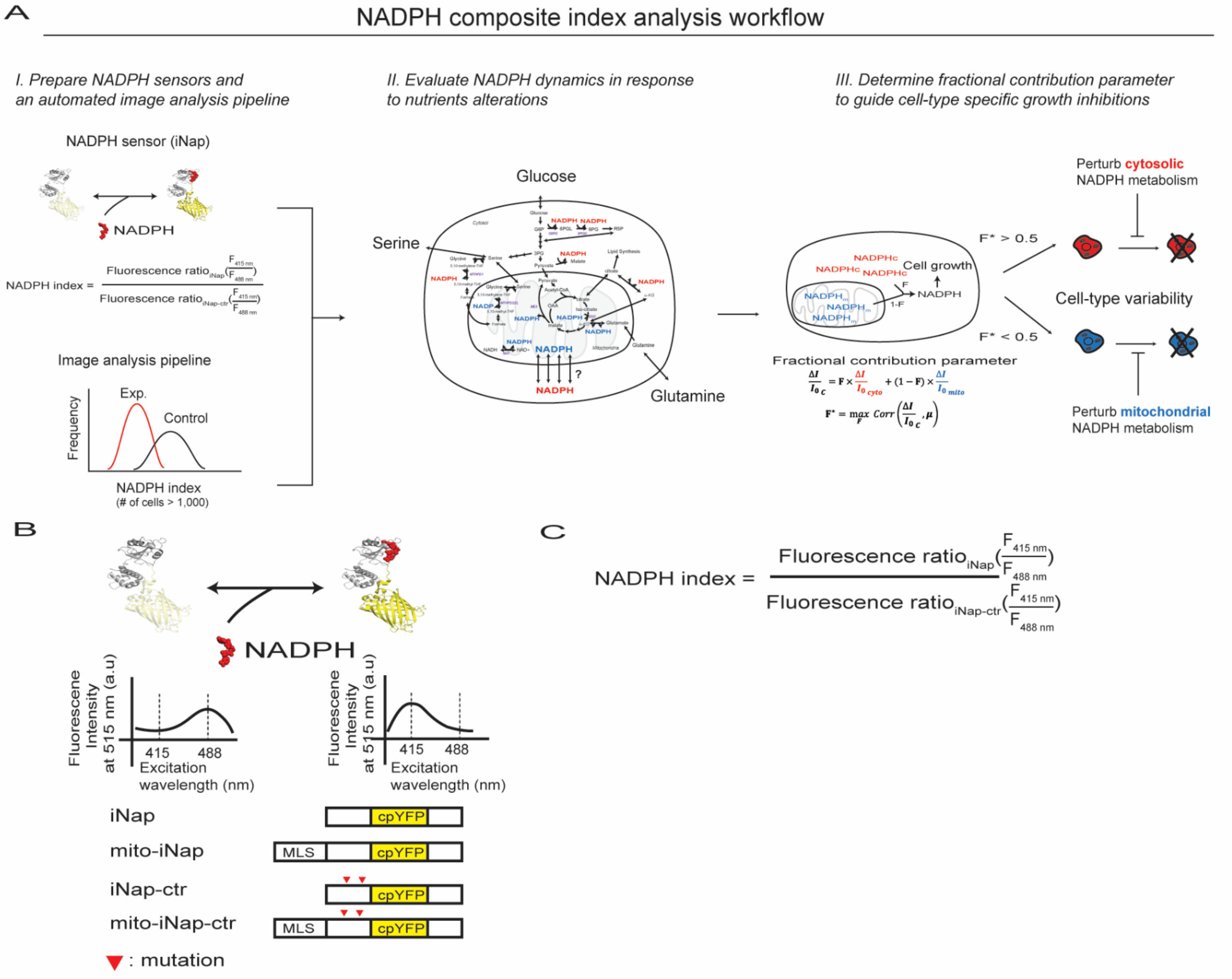
A
NADPH composite index analysis workflow
I. Prepare NADPH sensors and an automated image analysis pipeline
II. Evaluate NADPH dynamics in response to nutrients alterations
III. Determine fractional contribution parameter to guide cell-type specific growth inhibitions
NADPH sensor (iNap)
NADPH
Image analysis pipeline
Exp.
Control
Frequency
NADPH index
(# of cells > 1,000)
Glucose
Serine
Glutamine
NADPH
Cell growth
F* > 0.5
F* < 0.5
Cell-type variability
Perturb cytosolic NADPH metabolism
Perturb mitochondrial NADPH metabolism
Fractional contribution parameter
B
NADPH
Fluorescene Intensity at 515 nm (a.u)
415
488
Excitation wavelength (nm)
iNap
mito-iNap
iNap-ctr
mito-iNap-ctr
MLS
cpYFP
: mutation
C
NADPH index = Fluorescence ratio iNap (F 415 nm / F 488 nm) / Fluorescence ratio iNap-ctr (F 415 nm / F 488 nm)

Figure S1

**Figure S1, related to Figure 1.**

**(A)** Detailed NADPH composite index analysis workflow. It consists of 1) preparation of NADPH sensors and an automated image analysis pipeline, 2) evaluation of NADPH dynamics in response to varying extracellular nutrients, and 3) determination of fractional contribution parameter by correlating cytosolic and mitochondrial NADPH dynamics to growth rates in cancer cells. The parameter serves as a guide to target cytosolic or mitochondrial NADPH metabolism for selective cancer cell growth inhibitions. **(B)** Design of iNap constructs, including iNap, mito-iNap, iNap-ctr, and mito-iNap-ctr. The emission wavelength at 515 nm changes upon excitation at 415 and 488 nm change in presence of NADPH. NADPH index represents a pH-corrected sensor output as defined by the ratio of iNap fluorescence ratio to that of iNap-ctr. The red triangle represents the location of mutation. **(C)** Definition of NADPH index, which is the ratio between the fluorescence ratio of iNap sensor and that of iNap-ctr sensor.

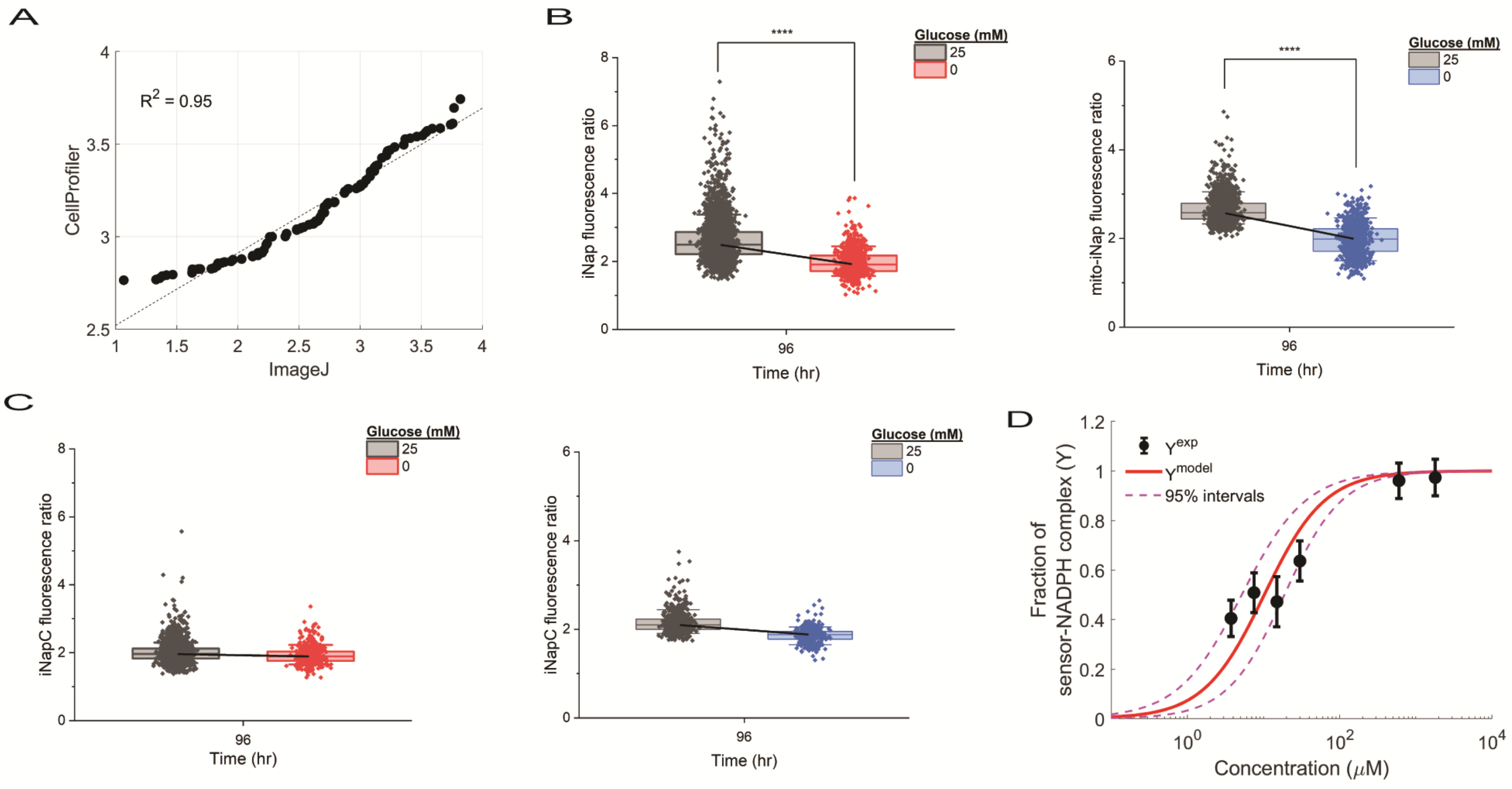
A
$R^2$ = 0.95
CellProfiler
ImageJ
B
Glucose (mM)
25
0
****
iNap fluorescence ratio
Time (hr)
96
mito-iNap fluorescence ratio
C
iNapC fluorescence ratio
D
Y^exp
Y^model
95% intervals
Fraction of sensor-NADPH complex (Y)
Concentration (μM)

Figure S2

### Figure S2, related to Figure 2.

**(A)** Comparison of the fluorescence ratio of iNap images based on our method and ImageJ, followed by Pearson's correlation analysis. **(B)** Fluorescence intensities with excitation at 415 and 488 nm of individual cells for iNap and mito-iNap in absence of glucose condition (n = 3916, 1561, 917 cells for iNap+Glc, iNap-Glc, mito-iNap-Glc, mito-iNap-Glc conditions). A total three biological replicates, each of which containing six technical replicates were used for analysis. **(C)** fluorescence ratio of iNapC and mito-iNapC was calculated and represented as a box plot (n = 2457, 500, 1561, 917 for iNapC+Glc, iNapC-Glc, mito-iNapC-Glc, mito-iNapC-Glc conditions). **(D)** Fraction of sensor-NADPH complex (Y) was measured experimentally ($Y^{\mathrm{exp}} = \frac{R' - R'_{min}}{R'_{max} - R'_{min}}$) and compared with the Y ($Y^{model} = \frac{S^n}{\mathrm{K_d} + S^n}$) calculated from a Hill–Langmuir equation. $R'$ represents the NADPH index, a normalized iNap fluorescence ratio to iNapC fluorescence ratio. $R'_{max}$ and $R'_{min}$ are maximum and minimum NADPH index. S represents the NADPH concentration and $K_d$ represents dissociation constants. 3.6 μM was used for $K_d$. n is the Hill coefficient, to which the experimental data was fitted. Goodness of fit was evaluated to provide 95% confidence interval ($n_{fit}$ = 1.089 with lower and upper bounds for 1.029 and 1.149).

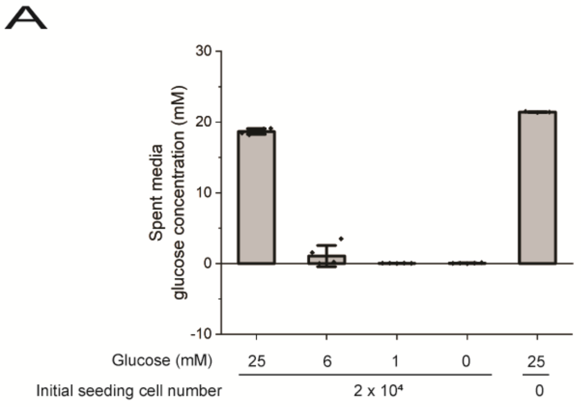
A
Spent media glucose concentration (mM)
30
20
10
0
-10
Glucose (mM)
25
6
1
0
25
Initial seeding cell number
2 x 10⁴
0

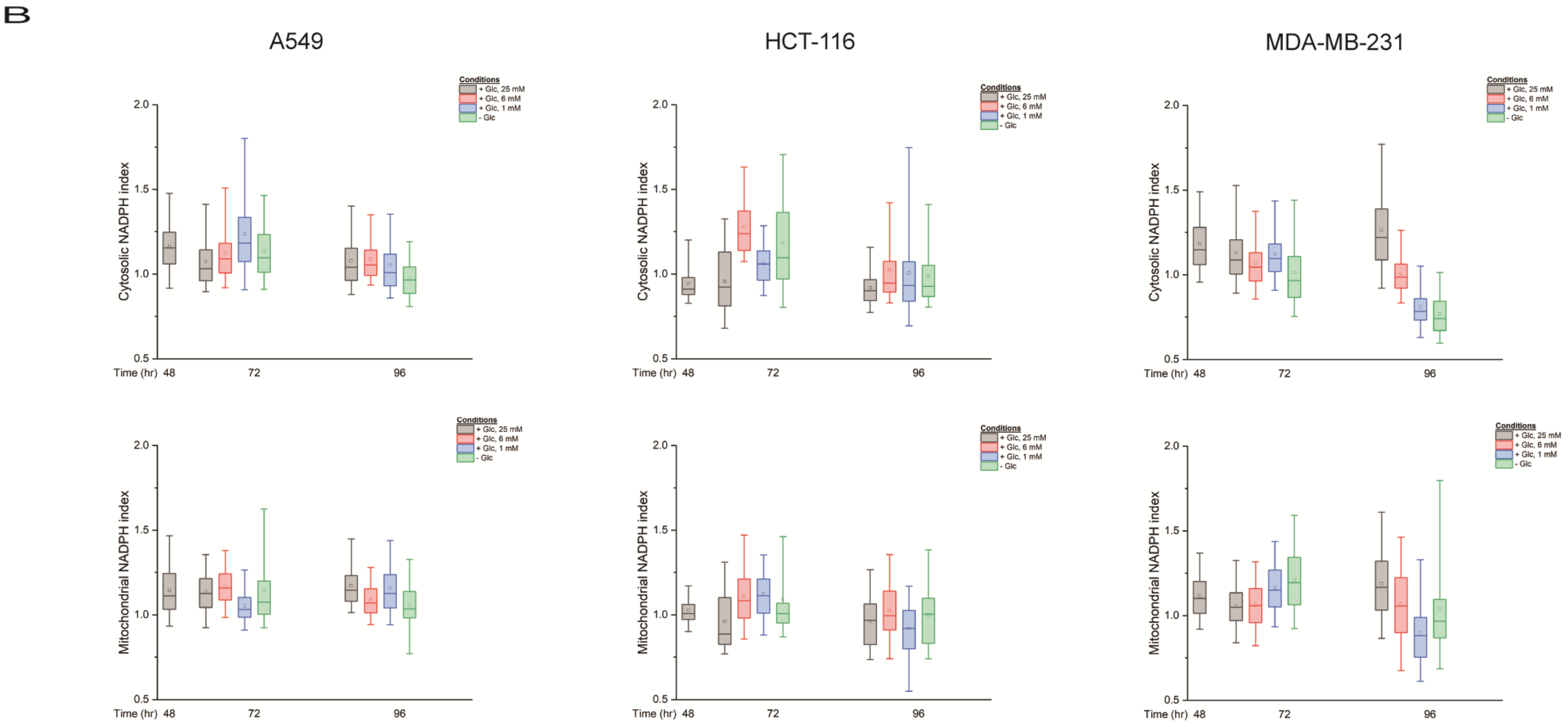
B
A549
HCT-116
MDA-MB-231
Conditions
+ Glc, 25 mM
+ Glc, 6 mM
+ Glc, 1 mM
- Glc
Cytosolic NADPH index
Mitochondrial NADPH index
2.0
1.5
1.0
0.5
Time (hr)
48
72
96

Figure S3

**Figure S3, related to Figure 3.**

**(A)** glucose concentration in the spent media was analyzed after 96 hours of culturing cells in DMEM with 10 % FBS with 25, 6, 1, and 0 mM glucose using Yellow Spring Instruments (YSI) 2950 instrument. **(B)** cytosolic and mitochondrial NADPH indices were measured after 48, 72, and 96 hours in A549, HCT-116, and MDA-MB-231 cell lines. Boxplot represents the $25^{th}$, $50^{th}$, and $75^{th}$ percentiles with whiskers indicating $5^{th}$ and $95^{th}$ percentile, in which the data were collected and combined from three biological replicates. For Hela, $n_{cyto}$ = 1323, 260, 253, 278, 3641, 1574, 1195, 1428, 7374, 3211, 2307, 1729, 9579, 4583, 2786, 2042, 14466, 5907, 2961, 1926; $n_{mito}$ = 739, 421, 694, 434, 1675, 1159, 1179, 963, 1694, 2157, 1662, 1587, 2901, 3380, 1752, 1979, 3341, 4390, 2365, 1947 were used. For A549, $n_{cyto}$ = 175, 407, 264, 256, 193, 376, 336, 280, 88; $n_{mito}$ = 196, 294, 277, 267, 161, 204, 236, 135, 40 were used. For HCT-116, $n_{cyto}$ = 95, 51, 53, 61, 36, 204, 109, 71, 198; $n_{mito}$ = 195, 146, 73, 105, 127, 94, 74, 30, 99. For MDA-MB-231, $n_{cyto}$ = 190, 662, 373, 312, 113, 5076, 1024, 318, 130; $n_{mito}$ = 312, 275, 326, 461, 302, 1104, 401, 147, 193 were used.

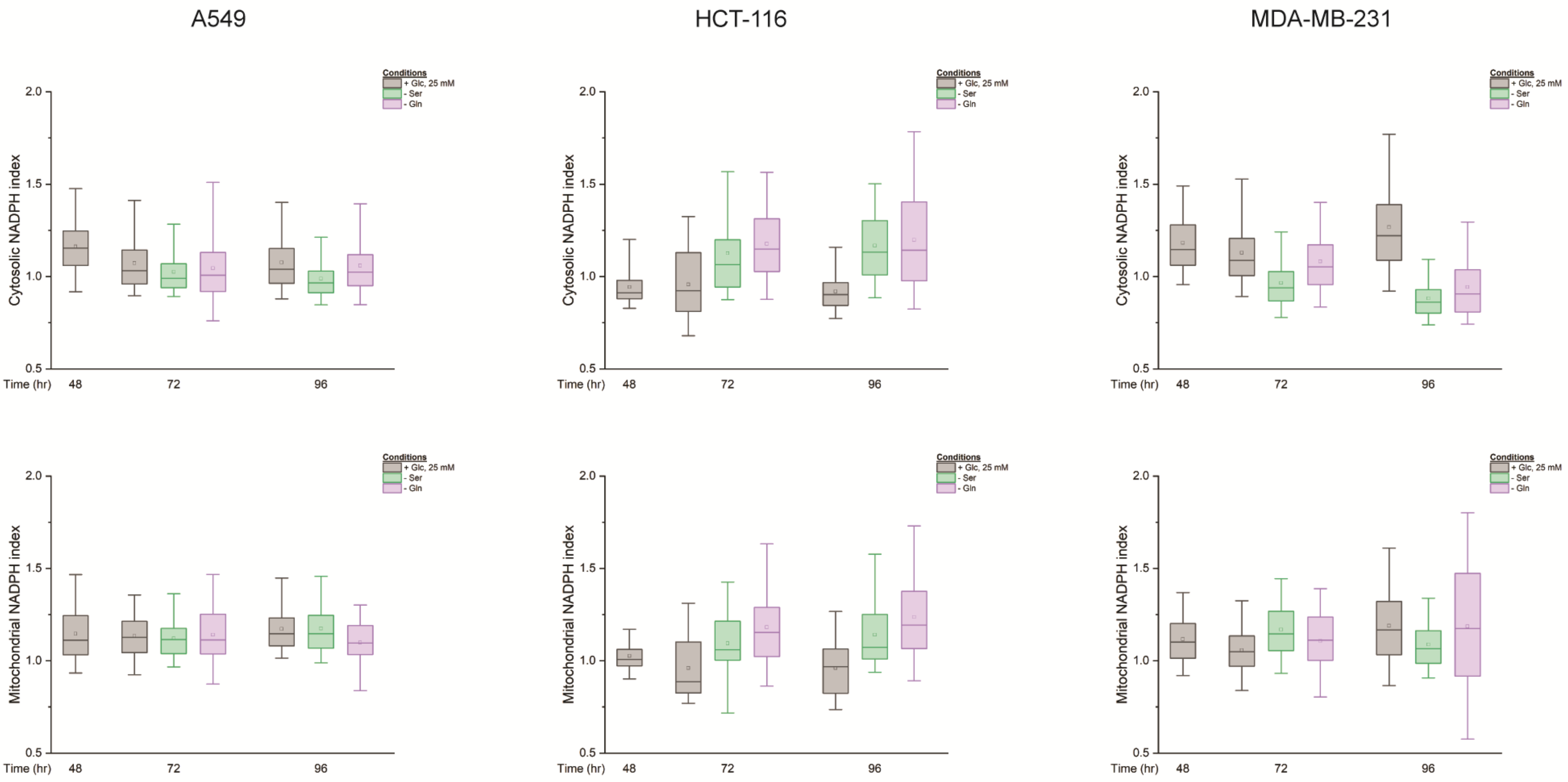
A549
HCT-116
MDA-MB-231
Conditions
+ Glc, 25 mM
- Ser
- Gln
Cytosolic NADPH index
Mitochondrial NADPH index
Time (hr)
48
72
96
2.0
1.5
1.0
0.5

**Figure S4**

**Figure S4, related to Figure 4.**

Cytosolic and mitochondrial NADPH indices were measured after 48, 72, and 96 hours in A549, HCT-116, and MDA-MB-231 cell lines. Boxplot represents the 25$^{th}$, 50$^{th}$, and 75$^{th}$ percentiles with whiskers indicating 5$^{th}$ and 95$^{th}$ percentile, in which the data were collected and combined from three biological replicates. For Hela, $n_{cyto,ser}$ = 2185, 2565, 4114, 3841, 4995, 4638, 8270, 5646; $n_{mito,ser}$ = 830, 981, 1824, 2098, 1871, 2322, 3260, 3437 were used for analysis. $n_{cyto,gln}$ = 2185, 2054, 4114, 3901, 4995, 2766, 8270, 4228; $n_{mito,gln}$ = 830, 895, 1824, 1965, 1871, 2081, 3260, 2164. For A549, $n_{cyto}$ = 175, 407, 167, 57, 376, 319, 82; $n_{mito}$ = 196, 294, 176, 66, 204, 214, 97 were used for analysis. For HCT-116, $n_{cyto}$ = 95, 51, 81, 44, 204, 182, 134; $n_{mito}$ = 195, 146, 57, 105, 94, 259, 135 were used for analysis. For MDA-MB-231, $n_{cyto}$ = 190, 662, 291, 155, 5076, 746, 284; $n_{mito}$ = 312, 275, 262, 298, 1104, 392, 308 were used for analysis.

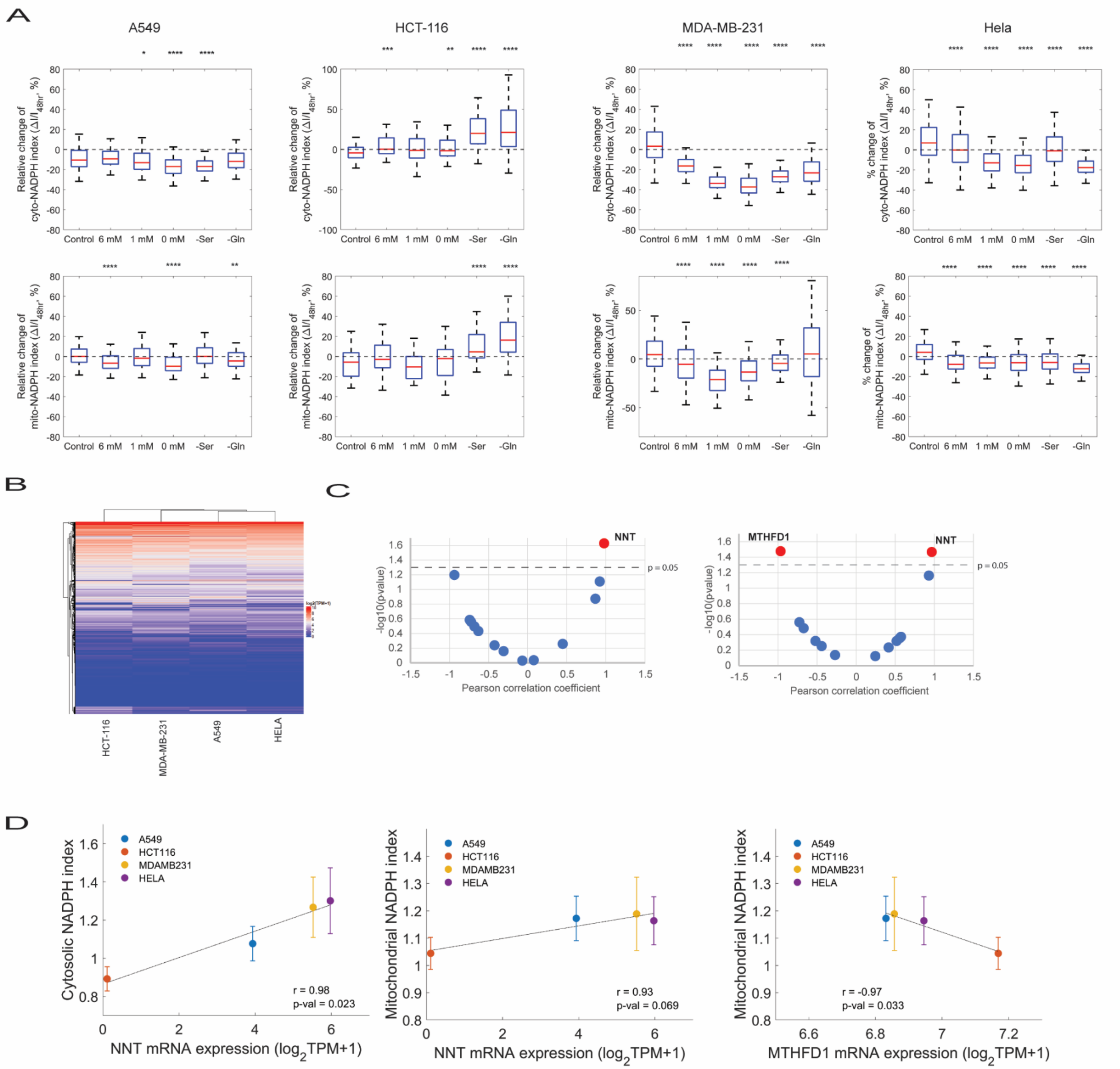
A
A549
HCT-116
MDA-MB-231
Hela
Relative change of cyto-NADPH index (ΔI/I48hr, %)
% change of cyto-NADPH index (ΔI/I48hr, %)
Control 6 mM 1 mM 0 mM -Ser -Gln
Relative change of mito-NADPH index (ΔI/I48hr, %)
% change of mito-NADPH index (ΔI/I48hr, %)
B
HCT-116
MDA-MB-231
A549
HELA
C
NNT
MTHFD1
p = 0.05
-log10(p-value)
Pearson correlation coefficient
D
A549
HCT116
MDAMB231
HELA
Cytosolic NADPH index
Mitochondrial NADPH index
r = 0.98
p-val = 0.023
r = 0.93
p-val = 0.069
r = -0.97
p-val = 0.033
NNT mRNA expression (log2TPM+1)
MTHFD1 mRNA expression (log2TPM+1)

**Figure S5**

### Figure S5, related to Figure 5

**(A)** relative change of cytosolic and mitochondrial NADPH indices (ΔI = $I_{96hr}$-$I_{48hr}$) to the indices at 48 hours across different nutrient conditions. Cells were cultured in DMEM with 25 mM glucose (control), varying glucose concentrations, and without serine or glutamine. **(B)** Hierarchical clustering analysis on 19,177 RNA-seq data on A549, HCT-116, Hela, and MDA-MB-231 (out of 1375 cell lines) was performed. **(C)** The relationship between cytosolic (left panel) or mitochondrial (right panel) NADPH indices and gene expression levels of NADPH generating enzymes in four cancer cell lines is evaluated through correlation analysis. Volcano plots represent the -$\log_{10}$(p-value) against person correlation coefficient. The dotted line represents a threshold p-value of 0.05. **(D)** Scatter plots representing cytosolic or mitochondrial NADPH indices against gene expression levels of either NNT and MTHFD1.

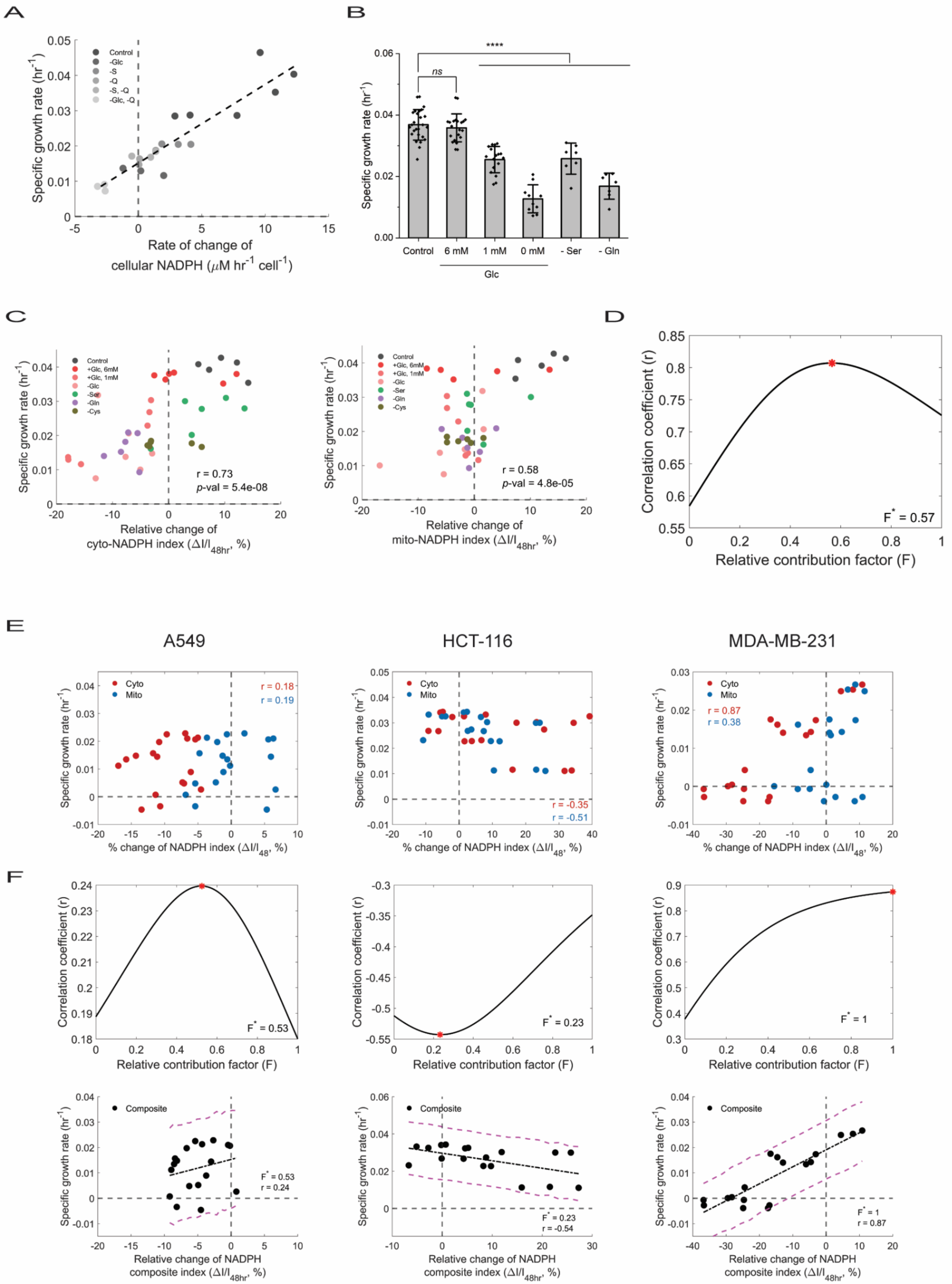
A
B
C
D
E
F
Specific growth rate (hr-1)
Rate of change of cellular NADPH (μM hr-1 cell-1)
Control
-Glc
-S
-Q
-S, -Q
-Glc, -Q
ns
****
6 mM
1 mM
0 mM
- Ser
- Gln
Glc
+Glc, 6mM
+Glc, 1mM
-Ser
-Gln
-Cys
r = 0.73
p-val = 5.4e-08
Relative change of cyto-NADPH index (ΔI/I48hr, %)
r = 0.58
p-val = 4.8e-05
Relative change of mito-NADPH index (ΔI/I48hr, %)
Correlation coefficient (r)
F* = 0.57
Relative contribution factor (F)
A549
HCT-116
MDA-MB-231
Cyto
Mito
r = 0.18
r = 0.19
r = -0.35
r = -0.51
r = 0.87
r = 0.38
% change of NADPH index (ΔI/I48, %)
F* = 0.53
F* = 0.23
F* = 1
Composite
r = 0.24
r = -0.54
Relative change of NADPH composite index (ΔI/I48hr, %)

**Figure S6**

### Figure S6, related to Figure 6

**(A)** Specific growth rate as a function of the rate of change of cellular NADPH. Each data represents values obtained from individual conditions described in the legends. **(B)** The growth rates of Hela cells were measured under different nutrient conditions and statistical significance was determined using t-tests, with p-values provided. P-values less than 0.05 were considered significant (*P<0.05, **P<0.01, ***P<0.001, ****P<0.0001). **(C)** The relationship between the growth rate of cells and the change in cytosolic or mitochondrial NADPH indices under varying nutrient conditions was evaluated using Pearson correlation coefficient and p-values. **(D)** The relationship between the NADPH composite index and growth rates varied depending on the fractional contribution parameter (F), as shown by the correlation coefficient (y-axis). **(E)** The correlation between growth rates and cytosolic (red) or mitochondrial (blue) NADPH indices was examined across various cancer cell lines. Statistical analysis was conducted using Pearson correlation coefficient (r) and p-values. **(F)** The optimal contribution parameter was determined and the growth rate was expressed as a function of the NADPH composite index with an optimized contribution parameter across cancer cell lines.

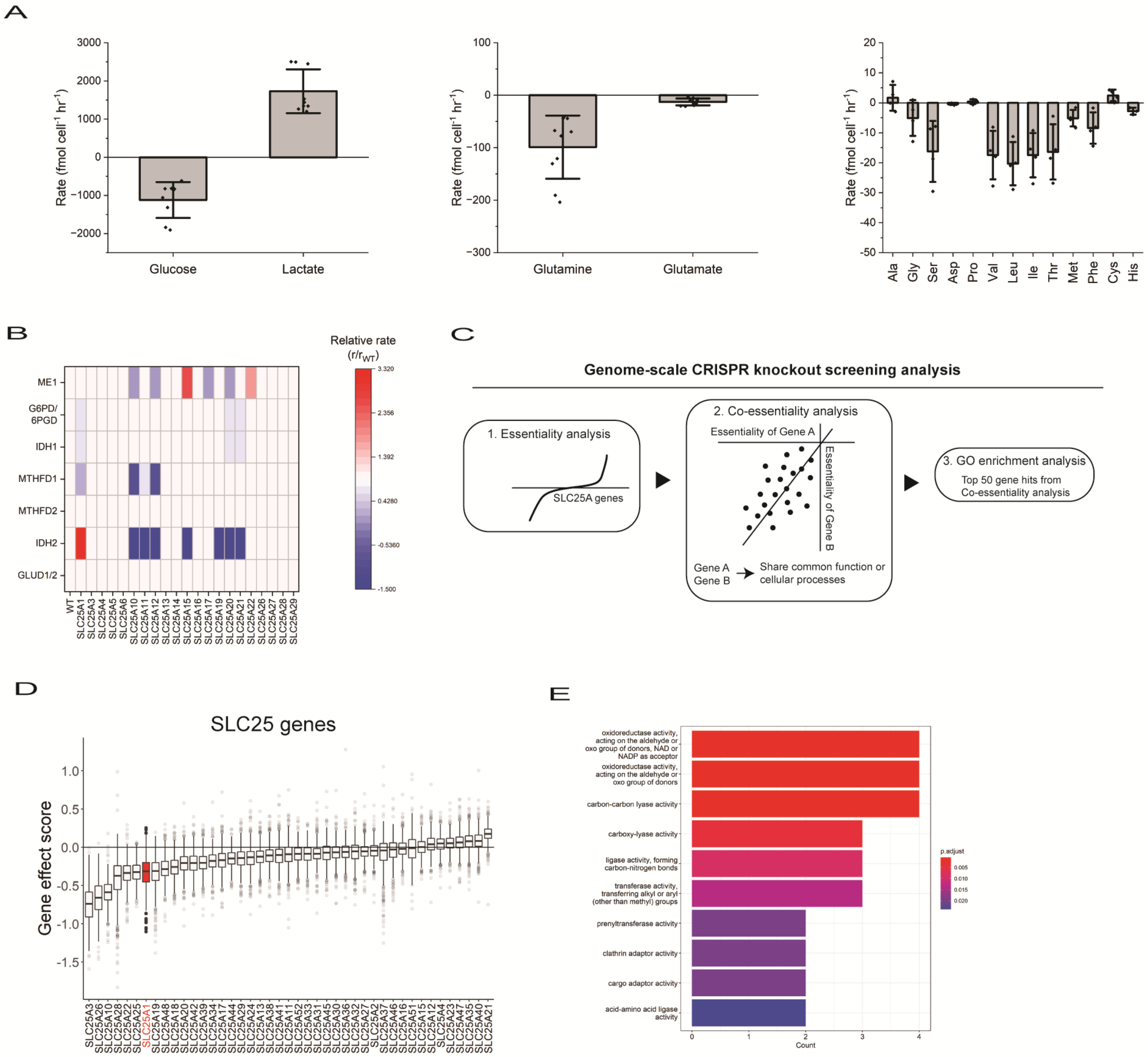
A
Rate (fmol cell-1 hr-1)
Glucose
Lactate
Glutamine
Glutamate
Ala
Gly
Ser
Asp
Pro
Val
Leu
Ile
Thr
Met
Phe
Cys
His
B
Relative rate (r/rWT)
ME1
G6PD/6PGD
IDH1
MTHFD1
MTHFD2
IDH2
GLUD1/2
WT
SLC25A1
SLC25A3
SLC25A4
SLC25A5
SLC25A6
SLC25A10
SLC25A11
SLC25A12
SLC25A13
SLC25A14
SLC25A15
SLC25A16
SLC25A17
SLC25A19
SLC25A20
SLC25A21
SLC25A22
SLC25A26
SLC25A27
SLC25A28
SLC25A29
3.320
2.356
1.392
0.4280
-0.5360
-1.500
C
Genome-scale CRISPR knockout screening analysis
1. Essentiality analysis
SLC25A genes
2. Co-essentiality analysis
Essentiality of Gene A
Essentiality of Gene B
Gene A
Gene B
Share common function or cellular processes
3. GO enrichment analysis
Top 50 gene hits from Co-essentiality analysis
D
SLC25 genes
Gene effect score
E
oxidoreductase activity, acting on the aldehyde or oxo group of donors, NAD or NADP as acceptor
oxidoreductase activity, acting on the aldehyde or oxo group of donors
carbon-carbon lyase activity
carboxy-lyase activity
ligase activity, forming carbon-nitrogen bonds
transferase activity, transferring alkyl or aryl (other than methyl) groups
prenyltransferase activity
clathrin adaptor activity
cargo adaptor activity
acid-amino acid ligase activity
Count
p.adjust

**Figure S7**

### Figure S7, related to Figure 7.

**(A)** Extracellular uptake or release rates of glucose (n = 9), lactate (n = 9), glutamate (n = 10), glutamine (n = 10), and other amino acids (n = 4). **(B)** Relative change of reaction rates of NADPH generating enzymes upon knockout of respective SLC25 genes. **(C)** Schematic representing the genome-scale CRISPR knockout screening analysis. **(D)** Gene-essentiality analysis on SLC25 genes across 739 cell lines. **(E)** Gene ontology enrichment analysis on the top fifty genes with the highest correlation scores from the co-essentiality analysis for SLC25A1. Molecular function mode was used for enrichment analysis.

**Table S1. Top 10 SLC25 genes with lowest gene effect score from Achilles Gene Effect database**

| Gene name | Gene effect Score | Aliases | Substrates |
|---|---|---|---|
| SLC25A3 | -0.74 | Phosphate carrier | Phosphate |
| SLC25A26 | -0.66 | S-adenosyl methionine carrier | S-adenosyl methionine, S-adenosyl homocysteine |
| SLC25A10 | -0.59 | Dicarboxylates carrier | Succinate, malate, phosphate, sulphate, thiosulphate |
| SLC25A28 | -0.37 | mitoferrin-2 | Unknown |
| SLC25A22 | -0.34 | Glutamate carrier 1 | Glutamate |
| SLC25A25 | -0.33 | ATP-MG/Pi carrier | ATP-Mg, Pi, ADP, ATP |
| SLC25A1 | -0.31 | Citrate carrier | Citrate, isocitrate, malate |
| SLC25A19 | -0.31 | Thiamine-pyrophosphate carrier | thiamine-pyrophosphate, thiamine-monophosphate, (deoxy)nucleotides |
| SLC25A48 | -0.29 | unknown | Unknown |
| SLC25A18 | -0.26 | Glutamate carrier 2 | Glutamate |

**Table S2. Pathway enrichment analysis for selected SLC25 genes* with corresponding top 50 co-essential genes.**

| Gene name | GO molecular function | Adjusted p-value | GO biological process | Adjusted p-value |
|---|---|---|---|---|
| SLC25A1 | oxidoreductase activity, acting on the aldehyde or oxo group of donors, NAD or NADP as acceptor | 4.9E-4 | nucleotide biosynthetic process | 2.29E-8 |
| SLC25A3 | oxidoreduction-driven active transmembrane transporter activity | 2.6E-37 | aerobic respiration | 1.5E-44 |
| SLC25A19 | electron transfer activity | 7.5E-26 | mitochondrial gene expression | 3.38E-15 |
| SLC25A26 | structural constituent of ribosome | 2.3E-24 | mitochondrial gene expression | 4.4E-42 |
| SLC25A28 | catalytic activity, acting on DNA | 0.036 | DNA-templated DNA replication | 1.66E-7 |
| SLC25A10 | No statistically significant results | | | |
| SLC25A18 | No statistically significant results | | | |
| SLC25A22 | No statistically significant results | | | |
| SLC25A25 | No statistically significant results | | | |
| SLC25A48 | No statistically significant results | | | |

* Top ten SLC25 genes with the lowest scores from the essentiality analysis were selected
